# Single-cell analysis reveals distinct fibroblast plasticity during tenocyte regeneration in zebrafish

**DOI:** 10.1101/2023.04.18.537403

**Authors:** Arsheen M. Rajan, Nicole L. Rosin, Elodie Labit, Jeff Biernaskie, Shan Liao, Peng Huang

## Abstract

Despite their importance in tissue maintenance and repair, fibroblast diversity and plasticity remain poorly understood. Using single-cell RNA sequencing, we uncover distinct sclerotome-derived fibroblast populations in zebrafish, including progenitor-like perivascular/interstitial fibroblasts, and specialized fibroblasts such as tenocytes. To determine fibroblast plasticity *in vivo*, we develop a laser-induced tendon ablation and regeneration model. Lineage tracing reveals that laser-ablated tenocytes are quickly regenerated by preexisting fibroblasts. By combining single-cell clonal analysis and live imaging, we demonstrate that perivascular/interstitial fibroblasts actively migrate to the injury site, where they proliferate and give rise to new tenocytes. By contrast, perivascular fibroblast-derived pericytes or specialized fibroblasts, including tenocytes, exhibit no regenerative plasticity. Interestingly, active Hedgehog (Hh) signaling is required for the proliferation of activated fibroblasts to ensure efficient tenocyte regeneration. Together, our work highlights the functional diversity of fibroblasts and establishes perivascular/interstitial fibroblasts as tenocyte progenitors that promote tendon regeneration in a Hh signaling-dependent manner.

**TEASER:** Perivascular/interstitial fibroblasts function as plastic progenitors during tenocyte regeneration.

## INTRODUCTION

Upon their initial discovery, fibroblasts are described as ‘spindle-shaped’ connective tissue cells with the ability to generate extracellular matrix (ECM) proteins (*1*, *2*). Since then, similar ECM-producing cells have been identified in almost all tissues and organs in vertebrates, and fibroblasts form one of the most abundant cell types in the body. Apart from depositing and remodeling ECM components, fibroblasts have been shown to play specialized roles in paracrine cell signaling, immune regulation, and stem cell niche maintenance (*3*, *4*). Furthermore, numerous studies have shown that fibroblasts can acquire an activated ‘myofibroblast’ state, upregulating matrix production and releasing cytokines to promote quick wound closure and regeneration upon tissue injury (*5–7*). Consequently, dysregulation of fibroblasts is often associated with connective tissue disorders and fibrotic scarring (*3*, *6*, *8*). However, despite their critical importance, the precise identity, behavior, and function of fibroblast populations *in vivo* remain poorly understood.

While fibroblasts have been extensively characterized in culture, the advent of single-cell RNA sequencing (scRNA-seq) has recently enabled in-depth exploration of fibroblast identity *in vivo*. Taking advantage of this, numerous studies have described inter- and intra-tissue heterogeneity of fibroblasts across several murine organs (*9–13*). In fact, fibroblasts from mouse skin, colon, heart, and skeletal muscles show vastly different ‘matrisomal’ (ECM-related) gene expression patterns and share less than 12% of their markers, suggesting that the transcriptional profile of fibroblasts is highly tailored to their specific tissue of residence (*12*). This idea is supported by an even broader study examining 16 mouse tissues which demonstrates that fibroblasts are not only heterogeneous but can be largely categorized into ‘specialized’ and ‘universal’ subpopulations (*13*). Specialized fibroblasts are restricted to single tissues and express tissue-specific markers, while universal fibroblasts are dispersed across multiple tissues, show elevated expression of ‘stemness’ genes, and are predicted to serve as progenitors for specialized fibroblasts (*13*). Interestingly, lineage tracing has shown that *Dpt^+^* universal fibroblasts become activated after subcutaneous tumor implantation, further supporting their ‘progenitor-like’ status *in vivo* (*13*). Orthologous injury-responsive fibroblast populations have also been identified in steady-state and perturbed human tissues, suggesting fibroblast diversity and specialization are conserved across species and disease states (*13–16*). These findings raise the exciting question of whether similar pools of ‘progenitor-like’ and ‘specialized’ fibroblasts are maintained in other vertebrates as well.

Zebrafish has emerged as a powerful model for studying tissue development and repair owing to its transparency, elevated regenerative capacity, and high synteny with humans (*17*, *18*). A few studies have characterized individual fibroblast populations in the trunks of zebrafish embryos. For instance, fibroblasts along the horizontal myosepta (*19*) and stromal reticular cells in the caudal vein plexus (*20*, *21*) have been shown to regulate lymphangiogenesis and hematopoiesis during zebrafish development, respectively. Similarly, tenocytes along the myotendinous junction (MTJ) (*22–25*), fin mesenchymal cells in the fin folds (*26*), and perivascular fibroblasts along intersegmental vessels (ISVs) (*27*) are known to provide structural support to their respective tissues of residence by depositing ECM. Several fibroblast populations have also been identified in single-cell profiling studies of the zebrafish trunk (*19*, *28*, *29*). However, no in-depth characterization of these populations has been performed to date and the full extent of fibroblast diversity and plasticity remains unknown.

We previously show that the sclerotome region of the somite gives rise to numerous fibroblast populations in the zebrafish trunk, including tenocytes, fin mesenchymal cells, interstitial fibroblasts, and several blood vessel-associated populations (*24*, *27*, *30*). In this study, we perform scRNA-seq to clarify that these sclerotome-derived fibroblasts comprise six subtypes with distinct transcriptional signatures and anatomical locations. Next, using a zebrafish model of tendon regeneration, we determine that these sclerotome-derived fibroblast subtypes show functional differences *in vivo*. Particularly, we demonstrate that ‘progenitor-like’ perivascular/interstitial fibroblasts, but not other fibroblast subtypes, actively migrate, proliferate, and differentiate into new tenocytes after tendon injury. Strikingly, this regenerative fibroblast response is mediated by the Hedgehog (Hh) signaling pathway. Altogether, our findings highlight transcriptional, spatial, and functional differences among fibroblast subpopulations in the zebrafish trunk and identify a role for perivascular/interstitial fibroblasts in tendon regeneration.

## RESULTS

### The sclerotome gives rise to fibroblasts in development

In our previous work, we established the homeobox transcription factor gene *nkx3-1* as a specific sclerotome marker in zebrafish (*24*, *30*). Using the *nkx3-1:Gal4; UAS:NTR-mCherry* transgenic line (*nkx3-1^NTR-mCherry^*, similar designations are used for all Gal4/UAS lines), we demonstrated that the sclerotome compartment of the somite gives rise to multiple fibroblast populations in the trunk, including tendon fibroblasts (tenocytes), fin mesenchymal cells in the fin fold, blood vessel-associated fibroblasts, and interstitial fibroblasts (*24*, *27*, *30*). To determine whether these fibroblast subtypes represent transcriptionally unique cell populations, we performed single-cell RNA sequencing (scRNA-seq). To enrich for sclerotome-derived fibroblasts, trunk sections of *nkx3-1^NTR-mCherry^*embryos at 52 hours post fertilization (hpf) were dissociated and mCherry^+^ cells were collected by fluorescence-activated cell sorting (FACS) (Figs. 1A, S1A). While sclerotome derivatives no longer actively express *nkx3-1* at 52 hpf, they continue to be labelled by mCherry due to perdurance of the protein and can be reliably isolated (Fig. S1A). A dataset of 2705 cells was generated after filtering for quality control metrics (Fig. S1B).

**Fig. 1.**
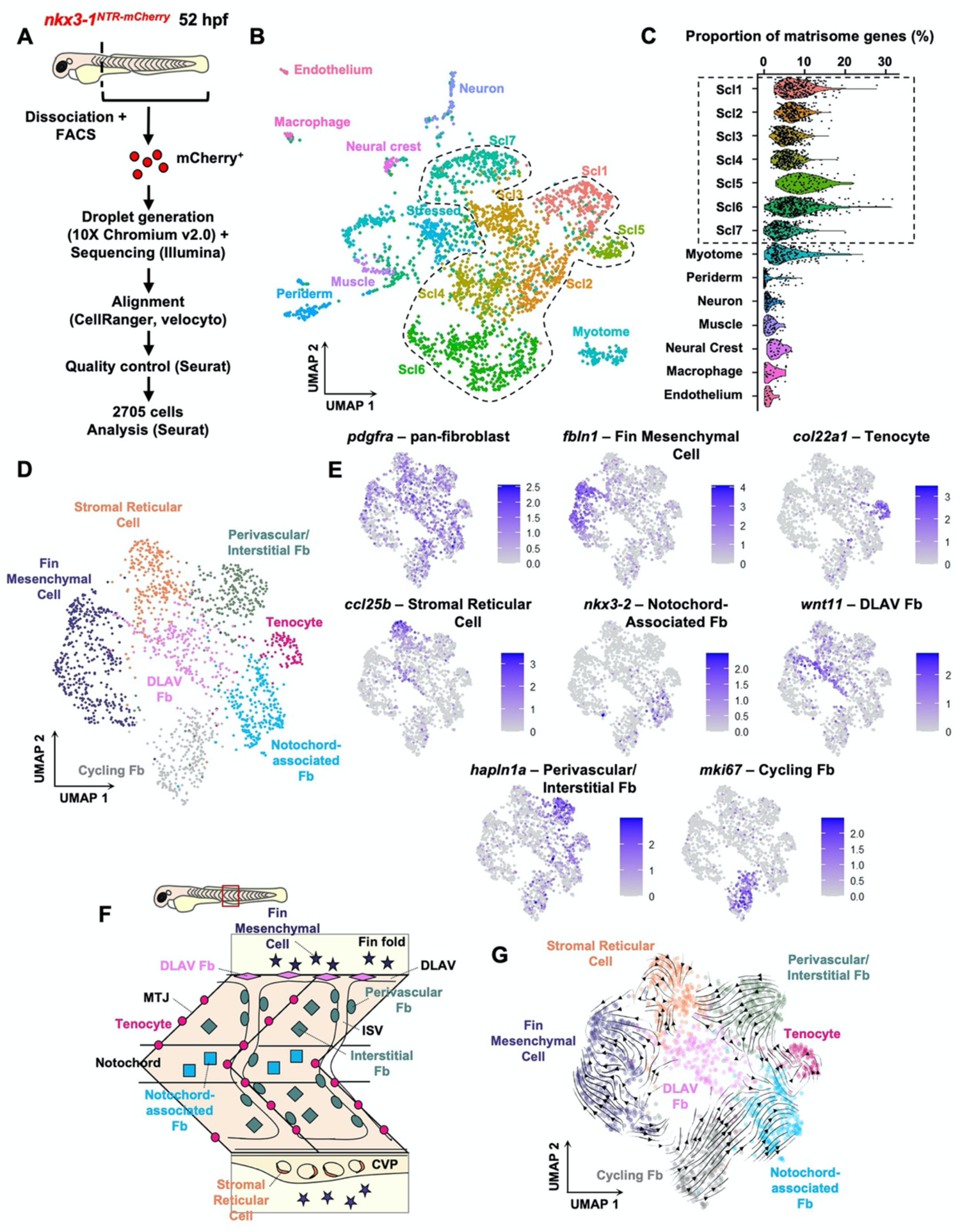
Single-cell RNA sequencing reveals heterogeneity and differential plasticity in sclerotome-derived fibroblasts. (**A**) Sample preparation pipeline for scRNA-seq of mCherry^+^ cells from 52 hpf *nkx3-1^NTR-mCherry^* zebrafish trunks. (**B**) UMAP projection of the post filtering scRNA-seq dataset. Dotted line indicates sclerotome-derived fibroblast clusters. (**C**) Proportion of ECM-related (matrisomal) transcripts detected per cell, grouped by cluster. Matrisome gene expression is enriched in sclerotome-derived fibroblast clusters (dotted line). (**D**) UMAP representation of subsetted fibroblast clusters from B, with distinct fibroblast subtypes labelled. (**E**) Feature plots showing expression of a pan-fibroblast marker, *pdgfra*, and one marker for each cluster identified in D. (**F**) Schematic depicting distribution of fibroblast subtypes from D in the zebrafish trunk, as determined in Figs. S4-6. (**G**) RNA velocity analysis of fibroblast clusters. Stream and arrowheads indicate direction of differentiation trajectory. Abbreviations: CVP, caudal vein plexus; DLAV, dorsal longitudinal anastomotic vessel; Fb, fibroblast; ISV, intersegmental vessel; MTJ, myotendinous junction; Scl, sclerotome.

UMAP projection of the filtered dataset produced 15 unique clusters, which were manually annotated according to expression of known markers (Figs. 1B, S2, Table S1). Based on expression of reported marker genes for the sclerotome (*postnb*, *twist1b*, *matn4*, *tgKi*) and its derivatives (*24*, *28*, *30*), we identified seven clusters comprising 1756 cells arising from the sclerotome: Scl1-7 (Figs. 1B, S3). Sclerotome-derived cell clusters (Scl1-7) were also enriched for extracellular matrix (ECM) genes and marked by classical pan-fibroblast markers (*pdgfra*, *col1a2*, *col5a1*, *dcn*) (*12*, *31*), confirming their identity as fibroblasts (Figs. 1C, S3A). It is worth noting that Scl1-7 clusters were not labelled by *nkx3-1*, consistent with *nkx3-1* no longer being expressed at this stage (Fig. S3B). In addition to sclerotome-derived fibroblasts, we also identified minor populations of muscle, myotome, and periderm populations, which have been previously observed in *nkx3-1^NTR-mCherry^* fish (*30*) (Figs. 1B, S2, Table S1). Additionally, small clusters of endothelial cells, neurons, and macrophages were also detected in our dataset (Figs. 1B, S2, Table S1). Since these populations did not represent fibroblasts, as evident by low concentration of matrisome genes and fibroblast markers (Figs. 1C, S3A), they were excluded from further analysis.

### Sclerotome-derived fibroblasts are heterogeneous

To examine sclerotome-derived fibroblasts more closely, relevant cells were subsetted and reanalyzed. Subsetted fibroblasts segregated into seven distinct subclusters (Fig. 1D). To determine the identity and spatial distribution of these fibroblast subpopulations, we examined the expression patterns of marker genes for each fibroblast subtype *in vivo*. Top markers for each cluster were selected using the following criteria: high expression in >25% of cells in cluster of interest, and >30% difference in proportion of expressing cells in cluster of interest versus all other clusters (Figs. 1E, S4A, Table S2). Three highly expressed markers for each cluster were then selected for validation. For each marker gene, in situ hybridization was performed in *nkx3-1^Kaede^* embryos at 52 hpf which were co-stained with anti-Kaede antibody to label all fibroblasts (Figs. S4-6).

Five fibroblast subclusters showed clear spatial restriction in their distribution (Fig. 1F). These included three known sclerotome-derived fibroblast populations: fin mesenchymal cells, tenocytes, and stromal reticular cells. As expected, the ‘fin mesenchymal cell’ cluster was labelled by genes previously identified in fin cells (*Kln1*, *and1*, *and2*; Table S2) (*26*, *30*, *32*, *33*) and known to be important for zebrafish fin development.

Consistently, all top markers examined for this cluster (*hgd*, *Kln1*, *hpdb*) were expressed in Kaede^+^ fibroblasts within the dorsal and ventral fin folds in *nkx3-1^Kaede^* embryos (Figs. 1E, S4A, S4B). Similarly, the ‘tenocyte’ cluster was labelled by known tenocyte genes (*scxa, cilp*; Table S2) (*22*, *30*), and all three top candidate markers tested (*col22a1*, *cilp*, *prelp*) showed specific expression in fibroblasts along the myotendinous junction (MTJ) (Figs. 1E, S4A, S4C). Previous studies have shown that stromal reticular cells arise from the sclerotome and become closely associated with the vasculature in the caudal vein plexus (CVP), where they play a critical role in the hematopoietic stem cell (HSC) development (*20*, *21*, *30*, *34*). Consistent with this, we identified a ‘stromal reticular cell’ (SRC) cluster within our dataset, with high-level marker expression (*clec19a*, *sfrp1a*, *ccl25b*) in CVP-associated fibroblasts (Figs. 1E, S4A, S5A). The remaining spatially restricted clusters did not match previously characterized fibroblast populations. Since marker genes for these clusters were enriched in cells surrounding the notochord (*nkx3-2*, *ccn2b*, *crispld1b*) and the dorsal longitudinal anastomotic vessel (DLAV) (*wnt11*, *zic1*, *ca6*), we termed these clusters ‘notochord-associated fibroblasts’ and ‘DLAV fibroblasts’, respectively (Figs. 1E, S4A, S5B, S5C).

The remaining two fibroblast clusters were more widely distributed across the zebrafish trunk (Fig. 1F). This included the ‘perivascular/interstitial fibroblast’ cluster, markers for which (*hapln1a*, *cdh11*, *pcdh18b*) labelled fibroblasts associated with the intersegmental vessels (ISVs), which we previously termed perivascular fibroblasts (*27*), and interstitial fibroblasts (*30*) located between the spinal cord and musculature (Figs. 1E, S4A, S6A). Further sub-clustering failed to resolve perivascular and interstitial fibroblasts from each other, suggesting that although these cells occupy distinct anatomical locations, they are transcriptionally indistinguishable at 52 hpf at our current sequencing resolution. Our final ‘cycling fibroblast’ cluster was enriched for genes commonly associated with cell cycle progression (*mki67*, *pcna*, *top2a*) (Figs. 1E, S4A). To visualize these proliferating fibroblasts *in vivo*, we stained for mitotic marker phospho-histone H3 (PHH3) (Fig. S6B), and separately performed an EdU pulsing assay from 52-53 hpf (Fig. S6C). In each experiment, pools of PHH3^+^/EdU^+^ cells comprised all the different fibroblast subtypes identified above (Figs. S6B-D). Therefore, cycling fibroblasts likely represent a mixture of fibroblast subtypes actively undergoing cell division, rather than a unique fibroblast subtype.

Since all the identified fibroblast subtypes are derived from the same embryonic source, we asked whether we could determine the lineage relationship amongst them. To do this, we performed RNA velocity inference on our projected dataset. All the spatially restricted fibroblast populations (tenocytes, fin mesenchymal cells, SRCs, DLAV fibroblasts, and notochord-associated fibroblasts) only showed intra-cluster velocity (Fig. 1G), suggesting they are terminally differentiated. By contrast, perivascular/interstitial fibroblasts showed notable velocity toward other clusters, with approximately half of the cells within this cluster transitioning toward the tenocyte or notochord-associated fibroblast fate and the other half toward the stromal reticular cell fate (Fig. 1G), suggesting that they may act as progenitors for other fibroblasts.

Altogether, our analysis identifies six major sclerotome-derived fibroblast subpopulations in the zebrafish trunk at 52 hpf, although we cannot rule out the existence of additional rare subpopulations that may not have been captured in our dataset due to low cell counts and read depths. Since our analysis focuses on an early embryonic stage, it is possible our defined fibroblast subtypes further differentiate, diverge into additional subpopulations, or are lost as development progresses.

### Zebrafish can regenerate trunk tenocytes after injury

Based on our RNA velocity results, we hypothesized that perivascular/interstitial fibroblasts function as progenitors for more specialized fibroblast subtypes such as tenocytes. To test this possibility *in vivo*, we developed a tendon regeneration model in zebrafish. Tenocytes in the dorsal region of a single myotendinous junction (MTJ) 1-2 somites anterior to the end of yolk extension were ablated with a high intensity laser (Fig. 2A). Laser ablation resulted in the localized loss of tenocytes and damaged muscle fibers centered around the laser focal point, while the dermal epithelium superficial to the injury site, neighboring ISVs, and underlying spinal cord and notochord structures remained largely intact (Fig. S7). In situ hybridization for tenocyte markers, *prelp* and *tnmd*, was performed on injured embryos from 1-4 days post injury (dpi) to track tenocyte regeneration (Figs. 2A-B). A large gap in staining was obvious along the injury site at 1 dpi, confirming that laser ablation resulted in localized tenocyte loss (Figs. 2B-F). By 2 dpi, this gap was significantly reduced and ectopic *prelp*/*tnmd* expression could be observed surrounding the injured MTJ (Figs. 2B-F). This was surprising as expression of these tenocyte markers is typically limited to the MTJ in uninjured fish. By 4 dpi, the lost tenocytes appeared to be completely regenerated and ectopic *prelp* expression around the MTJ was mostly lost (Figs. 2B-F).

**Fig. 2.**
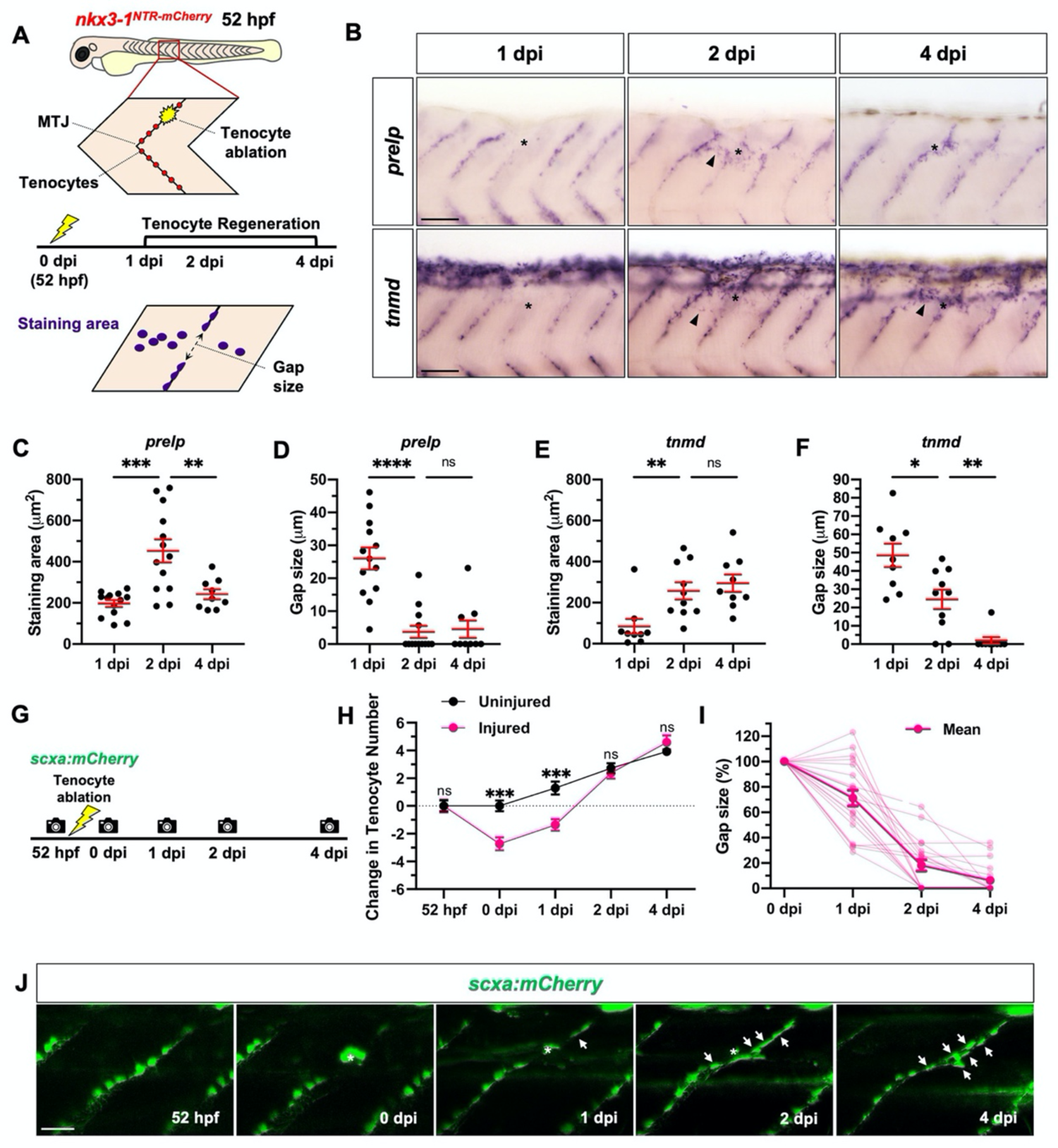
Zebrafish can regenerate trunk tenocytes after laser ablation. (**A**) Schematic representation of tenocyte ablation protocol and quantification shown in B-F. (**B**) In situ hybridization for tenocyte markers, *prelp* (top) and *tnmd* (bottom), at 1, 2, and 4 dpi showing increased gene expression (arrowheads) around the injury region (asterisks) at 2 dpi. *n* = 9-13 embryos/probe at each stage. (**C-F**) Quantification of staining area and gap size in stained embryos as described in A. (**G**) Experimental protocol to track tenocyte regeneration in live *scxa:mCherry* embryos. (**H**) Change in tenocyte number from 52 hpf to 4 dpi along injured and uninjured myotendinous junctions. (**I**) Quantification of gap between adjacent tenocytes at the injury site in each embryo from 0-4 dpi. (**J**) Representative images of one injured embryo from 52 hpf to 4 dpi, with several newly regenerated tenocytes (arrows) arising at the injured MTJ. Autofluorescent scar from laser ablation indicated by asterisks. *n* = 14 uninjured and 19 injured MTJs from 19 embryos. All Data shown as mean ± SEM. Statistics: Mann-Whitney *U*-test (C-F); Sidak’s multiple comparisons (H). Significance: p-value >0.05 (ns), <0.05 (*), <0.01 (**), <0.001 (***), <0.0001 (****). Scale bars: 50 μm (B); 25 μm (J).

We observed a similar timeline of tenocyte regeneration in live embryos using a tenocyte-specific transgenic reporter (*scxa:mCherry*) (*35*) (Figs. 2G-J). To track tenocyte regeneration, individual embryos were followed from 52 hpf to 4 days after laser ablation and tenocyte number was manually recorded at each stage (Fig. 2G). Laser ablation resulted in a mean loss of 3 tenocytes at injured MTJs at 0 dpi (Fig. 2H). Along uninjured MTJs, tenocyte number gradually increased by an average of 4 new tenocytes per MTJ from 52 hpf to 6 days post fertilization (dpf), suggesting a modest level of tenocyte development is still occurring at these stages (Fig. 2H). By contrast, the rate of tenocyte production was dramatically increased along injured MTJs in the same embryos, with an average of 7 new tenocytes being produced per MTJ from 0 to 4 dpi, suggesting that laser-induced MTJ injury promotes tenocyte proliferation and/or differentiation at the injury site (Figs. 2H,J). In fact, tenocyte number at injured MTJs was completely restored by 2 dpi (Fig. 2H). Tenocyte expansion was also accompanied by a concurrent reduction in gap size between adjacent tenocytes at injured MTJs (Figs. 2I-J).

To test whether MTJ architecture can also be restored after injury, we performed immunostaining for two MTJ markers, Vinculin and Thrombospondin 4b (Thbs4b) (*23*) (Fig. S8). Tenocyte ablation produced a clear gap in Vinculin distribution in 93% embryos (13/14) at 0 dpi (Figs. S8A,C,E). Similarly, 67% embryos (10/15) showed a gap in Thbs4b staining at 0 dpi (Figs. S8A,B,D). By 4 dpi, however, only 25% (4/16) and 10% (2/20) embryos still possessed gaps in Vinculin and Thbs4b staining, respectively (Fig. S8). Together, we conclude that majority of zebrafish embryos can efficiently regenerate lost tenocytes and the damaged MTJ within 4 days after laser-induced injury.

### New tenocytes are regenerated from preexisting sclerotome-derived fibroblasts

Next, we sought to determine the cellular origin of new tenocytes at the injury site. To test if new tenocytes were derived from neighboring fibroblasts, we utilized a photoconversion based lineage tracing approach (Fig. 3A). Briefly, we used the *col1a2^Kaede^* line to drive expression of photoconvertible Kaede protein in fibroblasts, which can be switched from green to red fluorescence upon exposure to UV light (*36*). At 52 hpf, the Kaede^green^ fluorescence in a two-somite region was photoconverted to label all preexisting fibroblasts in this area with Kaede^red^ (Fig. 3A). A single MTJ in the converted region was subsequently injured and embryos were followed from 0-4 dpi (Fig. 3A). Kaede^red^ fibroblasts could be seen populating the injury site as early as 1 dpi (Fig. 3B). By 4 dpi, 90% of injured embryos (28/31) possessed Kaede^red^ fibroblasts along the regenerating MTJ (Fig. 3B). Importantly, these Kaede^red^ fibroblasts displayed the distinct tenocyte-like morphology (*23*, *24*) by 4 dpi, with small, rounded cell bodies and numerous long processes extending laterally along the regenerated MTJ (Figs. 3B,C). Indeed, 80% of these tenocyte-like Kaede^red^ fibroblasts (51/64) at the regenerated MTJ expressed *tnmd*, confirming their tenocyte identity (Fig. 3D). Consistent with this, we observed *prelp-* expressing fibroblasts around the injury site in 65% of 1 dpi embryos (11/17) and 93% of 2 dpi embryos (13/14), suggesting that most injury-responsive fibroblasts have started to differentiate into tenocytes at these stages (Fig. S9).

**Fig. 3.**
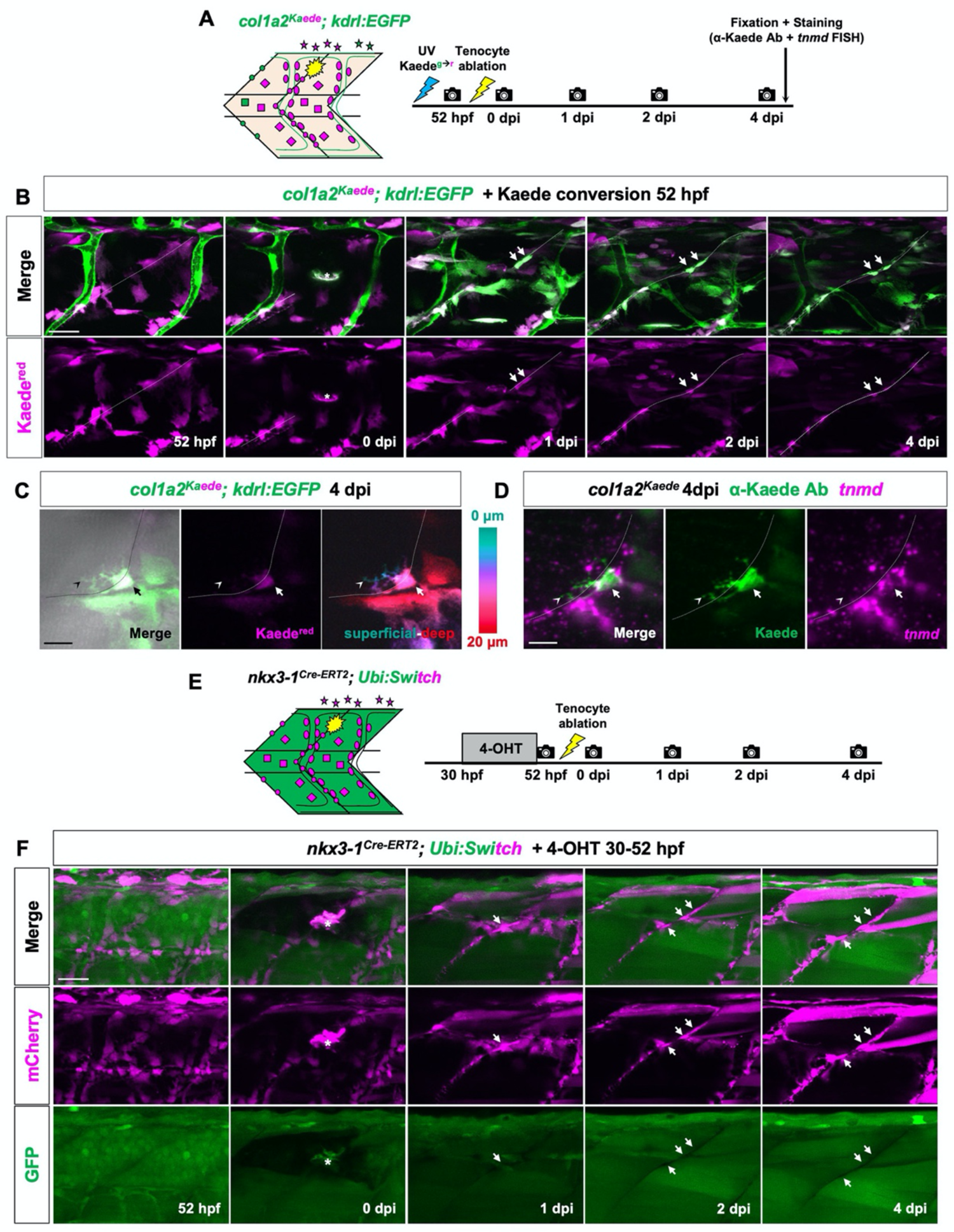
Sclerotome-derived fibroblasts give rise to new tenocytes after laser ablation. (**A**) Experimental timeline for fibroblast photoconversion and lineage tracing in *col1a2^Kaede^; kdrl:EGFP* embryos shown in B-D. (**B**) Representative images of one injured embryo from 52 hpf to 4 dpi. New Kaede^red^-positive fibroblasts (arrows) can be seen along the regenerating MTJ (dotted lines) from 1-4 dpi. *n* = 31 embryos. (**C**) Close up view of a single Kaede^red^-positive fibroblast (arrows) at a regenerated MTJ (dotted lines) at 4 dpi. Depth projection (right) reveals several tenocyte-like processes (notched arrowheads) extending laterally from the cell. (**D**) Co-immunostaining and fluorescent mRNA in situ hybridization showing tenocyte marker *tnmd* expression (magenta) in the traced fibroblast (arrows) from C. *n* = 51/64 cells from 21 embryos. Notched arrowheads indicate cell processes. (**E**) Protocol for Cre-mediated lineage tracing of sclerotome-derived fibroblasts after tenocyte ablation. (**F**) Representative images of a single injured embryo from 52 hpf to 4 dpi showing new mCherry^+^ tenocytes (arrows) at the regenerated MTJ. *n* = 12 embryos. Injury site is indicated by asterisks for all images. Scale bars: 25 μm (B,F); 10 μm (C,D).

As a complementary approach, we performed Cre-mediated lineage tracing of sclerotome-derived fibroblasts. Briefly, *nkx3-1:Gal4; UAS:Cre-ERT2; Ubi:Switch* embryos were treated with 4-hydroxy tamoxifen (4-OHT) at 30-52 hpf to permanently label sclerotome-derived fibroblasts with mCherry, immediately prior to tenocyte ablation (Fig. 3E). Consistent with cell tracing by Kaede conversion, 83% of injured embryos (10/12) possessed new mCherry^+^ tenocytes at the regenerated MTJ by 4 dpi (Fig. 3F). Together, our lineage tracing experiments demonstrate that preexisting sclerotome-derived fibroblasts give rise to new tenocytes after laser injury.

### Perivascular fibroblasts but not pericytes can differentiate into new tenocytes after injury

Our results demonstrate that sclerotome-derived fibroblasts are the primary source of new tenocytes after injury. This is consistent with our RNA velocity analysis which predicted perivascular/interstitial fibroblasts function as tenocyte progenitors during normal development (Fig. 1G). To directly test the plasticity of perivascular fibroblasts in response to tenocyte injury, we performed single-cell clonal analysis (Fig. 4A) by leveraging a highly mosaic *col1a2^Kaede^* transgenic line, which likely results from transcriptional silencing of UAS repeats over many generations (*37*). We photoconverted a single fibroblast from Kaede^green^ to Kaede^red^ before injuring the nearest MTJ (Fig. 4A). Injured embryos were imaged repeatedly from 0 to 4 dpi. Response of photoconverted cells was quantified as depicted by recording three primary metrics: cell behavior, clone composition, and clone size (Fig. 4B).

**Fig. 4.**
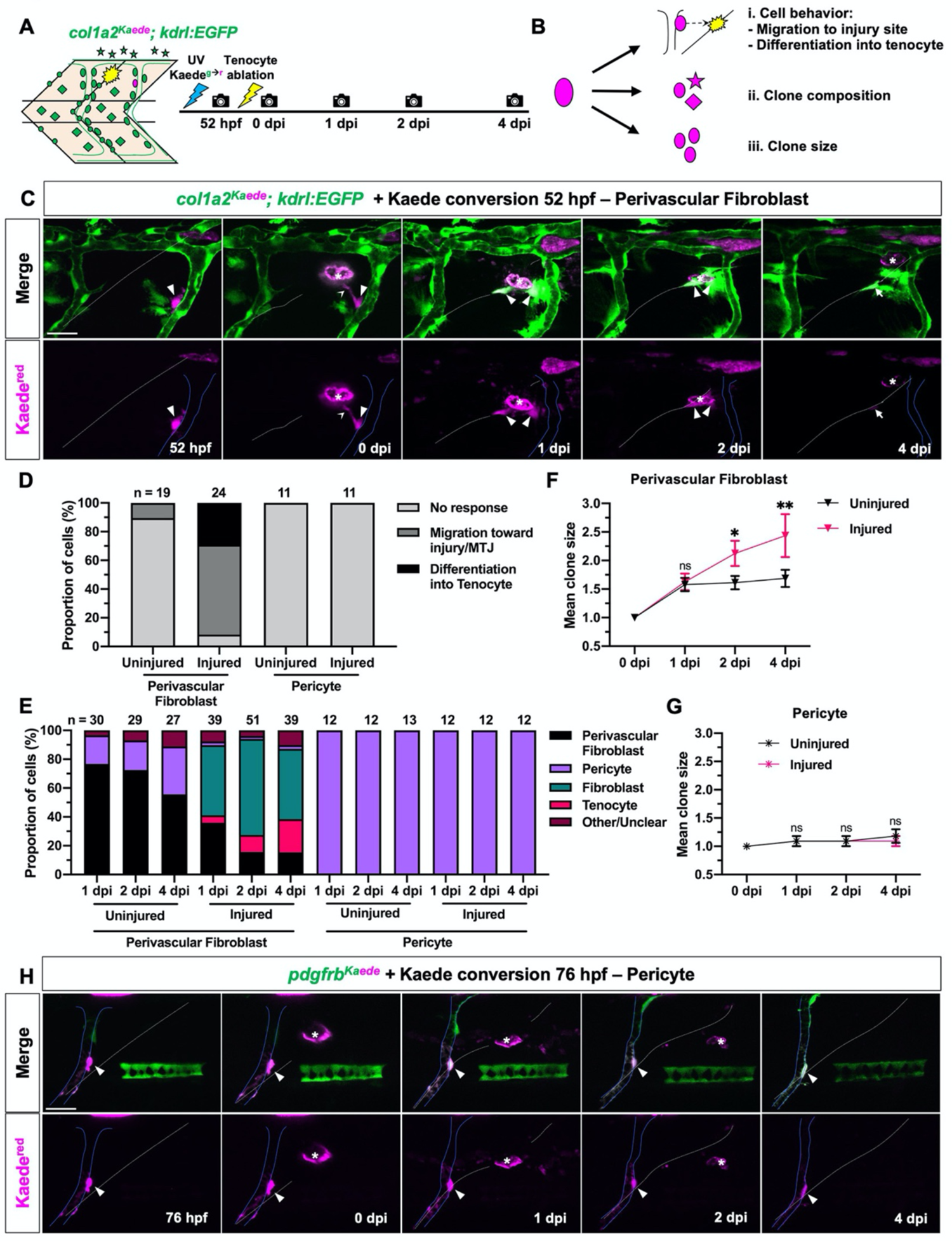
Perivascular fibroblasts but not pericytes respond to tendon injury. (**A**) Experimental protocol for clonal analysis of fibroblasts in *col1a2^Kaede^; kdrl:EGFP* embryos. A single fibroblast was converted from Kaede^green^ to Kaede^red^ fluorescence prior to tenocyte ablation, and injured fish were followed from 52 hpf to 4 dpi. (**B**) The three parameters quantified to describe the behavior of traced cells. (**C**) Representative images of a photoconverted perivascular fibroblasts from 52 hpf to 4 dpi. The traced cell (magenta, arrowheads) can be seen extending processes (notched arrowheads), migrating away from the ISV (solid blue lines), and giving rise to a new tenocyte (arrows) at the regenerating MTJ (white dotted lines) in response to tenocyte ablation. (**D**) Graph summarizing overall response of photoconverted perivascular fibroblasts and pericytes in injured and uninjured embryos. *n* = 19 (uninjured) and 24 (injured) perivascular fibroblasts; 11 (uninjured) and 11 (injured) pericytes. (**E**) Fibroblast subtype composition of the total clonal population across all traced perivascular fibroblasts and pericytes in injured and uninjured embryos at 1-4 dpi. Total clonal population size at each stage indicated above the graph. (**F-G**) Mean clone size of traced perivascular fibroblasts (F) and pericytes (G) from 0-4 dpi. (**H**) Representative images of a single photoconverted pericyte (magenta, arrowheads) from 76 hpf to 4 dpi in injured *pdgfrb^Kaede^* embryos. ISV and injured MTJ denoted by solid blue and dotted white lines, respectively. Data in F,G represented as mean ± SEM. Statistics: Sidak’s multiple comparisons (F,G). Significance: p-value >0.05 (ns), <0.05 (*), <0.01 (**). Scale bars: 25 μm.

In uninjured embryos, most perivascular fibroblasts (89%, 17/19) remained associated with the vasculature between 2-6 dpf, and never differentiated into tenocytes (Fig. 4D). In contrast, after tenocyte ablation, perivascular fibroblasts could be seen immediately extending projections toward the injury site, and by 1 dpi, most were no longer associated with the ISV (Fig. 4C). This dynamic process was captured by time-lapse imaging, where activated perivascular fibroblasts quickly migrated away from the ISV and underwent cell division at the injury site (Movie S1). By 4 dpi, 92% of traced perivascular fibroblasts (22/24) had migrated to the injury site (Figs. 4C,D). Of all traced cells, 29% (7/24) produced tenocyte-like progeny at 4 dpi which were localized to the regenerating MTJ extending several long processes laterally (Figs. 4C, 4D, S10A). Indeed, 95% of such tenocyte-like cells (21/22) were labelled by tenocyte marker, *col22a1*, confirming that perivascular fibroblasts differentiate into tenocytes after tendon injury (Fig. S10A). Interestingly, we also found 30% of other perivascular fibroblast-derived cells (6/20) that did not possess tenocyte-like morphology and were thus not categorized as tenocytes in our analysis, also expressed *col22a1*, suggesting that they are maturing tenocytes (Fig. S10A). Consistent with these results, clonal populations of traced perivascular fibroblasts in injured embryos were primarily comprised of activated fibroblasts (49%, 19/39) and newly generated tenocytes (23%, 9/39) at 4 dpi (Fig. 4E). Meanwhile, clonal populations in uninjured controls at the comparable stage (150 hpf) consisted primarily of ISV-associated perivascular fibroblasts (56%, 15/27) and pericytes (33%, 9/27) (Figs. 4E, S10B), consistent with our previous work showing perivascular fibroblasts give rise to pericytes during normal development (*27*). Notably, while both perivascular fibroblasts and pericytes are associated with ISVs, we previously demonstrated that they show distinct morphological characteristics (*27*). We therefore classified clonal progeny displaying long processes and elongated cell bodies as pericytes, and those with larger, globular cell bodies as perivascular fibroblasts (Fig. S10B) (*27*). Lastly, quantification revealed that the mean clonal population size of traced cells was also significantly increased in injured fish compared to uninjured controls at 2-4 dpi, suggesting tenocyte ablation induces perivascular fibroblast proliferation (Fig. 4F). Together, our clonal analysis demonstrates that perivascular fibroblasts function as tenocyte progenitors after tendon injury.

Since ISV-associated pericytes are derived from perivascular fibroblasts (*27*), we asked whether pericytes maintain the similar plasticity in response to tenocyte ablation. While *pdgfrb* is expressed in both perivascular fibroblasts (weak expression) and pericytes (strong expression), we have previously shown that our *pdgfrb:Gal4FF* line only drives expression in *pdgfrb^high^* pericytes in the embryonic zebrafish trunk (*27*, *38*). Using this pericyte-specific *pdgfrb^Kaede^* (*pdgfrb:Gal4FF; UAS:Kaede*) reporter, we performed single cell clonal analysis of pericytes. Since pericytes first appear around ISVs only at 3 dpf, embryos were photoconverted and injured at 76 hpf. Surprisingly, in contrast to perivascular fibroblasts, 100% of pericytes (11/11) remained tightly associated with the vasculature post injury and showed no migratory, differentiation or proliferative response (Figs. 4D,E,G,H). Furthermore, converted cells maintained their *pdgfrb* expression and characteristic long processes encircling the ISV, indicating that they retained their pericyte identity (Fig. 4H). In summary, we establish that perivascular fibroblasts are actively recruited to the injury site after tenocyte ablation where they contribute to tenocyte regeneration. However, this regenerative response is lost upon further differentiation to pericytes.

### Interstitial fibroblasts contribute to new tenocytes during development and regeneration

Our scRNA-seq analysis showed that interstitial fibroblasts, which are normally interspersed between the spinal cord and muscles (*30*), are transcriptionally indistinguishable from perivascular fibroblasts (Figs. 1D, S6A). Therefore, we hypothesized that interstitial and perivascular fibroblasts show similar response to tendon injury. As expected, interstitial fibroblasts were observed migrating to the injury site (75%, 12/16), undergoing morphological changes, and giving rise to new tenocytes post tenocyte ablation (38%, 6/16) between 0-4 dpi (Figs. 5A,C,D, S10C). All traced progeny identified as tenocytes were labelled by the tenocyte marker *col22a1* (Fig. S10C). This was supported by a corresponding increase in interstitial fibroblast clone size at 2 dpi in injured compared to uninjured embryos (Fig. 5E). Surprisingly however, unlike perivascular fibroblasts, 24% of traced interstitial fibroblasts (5/21) differentiated into tenocytes even in uninjured embryos (Figs. 5C-D). Therefore, even though perivascular and interstitial fibroblasts appear to be transcriptionally similar, here we find key functional differences between the two populations. Interstitial fibroblasts function as progenitors for tenocytes during normal development as well as post injury, whereas perivascular fibroblasts only do so upon injury. It is possible that while similar, interstitial and perivascular fibroblasts might possess subtle differences in transcriptional signature not picked up by our scRNA-seq experiment, explaining their difference in behavior in uninjured embryos.

**Fig. 5.**
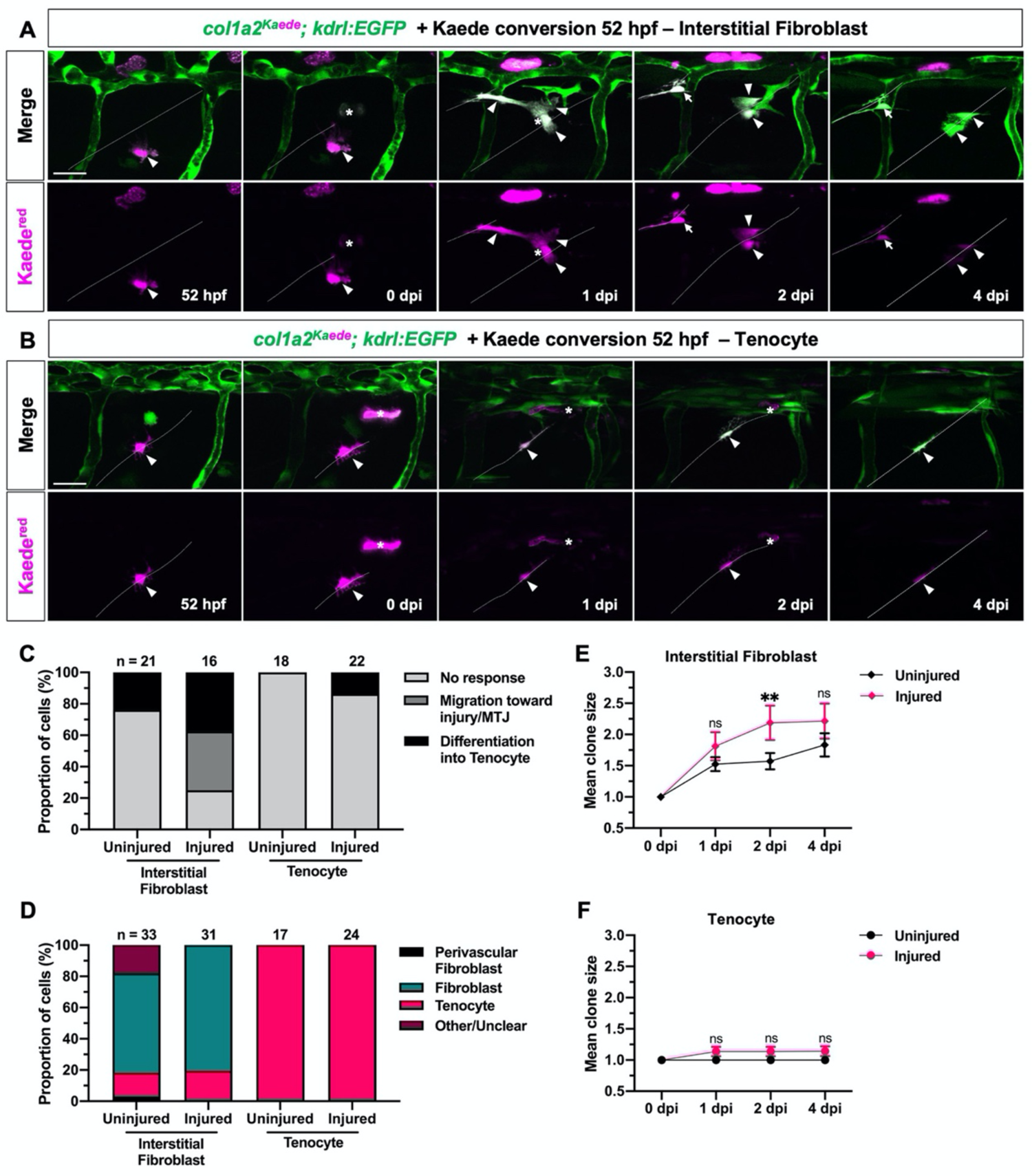
Interstitial fibroblasts but not tenocytes respond to tendon injury. (**A,B**) Representative images from clonal analysis of a single interstitial fibroblast (A) and tenocyte (B) in injured *col1a2^Kaede^; kdrl:EGFP* embryos, as described in Fig. 4A. The traced cells and their progeny (magenta) are indicated by arrowheads and the new tenocyte is denoted by arrows. Dotted lines label MTJs and injury site is denoted by asterisks in all images. (**C-F**) Quantification of interstitial fibroblast and tenocyte behavior, as depicted in Fig. 4B, showing overall response (C), clonal composition (D), and mean clone size (E,F) in injured and uninjured embryos. Note that the ‘differentiation into tenocyte’ category for tenocyte clonal analysis graph in C represents rare cases where traced tenocytes underwent cell division to give rise to new tenocytes at the injured MTJ. *n* = 21 (uninjured) and 16 (injured) interstitial fibroblasts; 18 (uninjured) and 22 (injured) tenocytes. Data in E,F represented as mean ± SEM. Statistics: Sidak’s multiple comparisons (E,F). Significance: p-value >0.05 (ns), <0.01 (**). Scale bars: 25 μm.

### Specialized sclerotome-derived fibroblasts do not respond to tendon injury

Unlike perivascular and interstitial fibroblasts, other sclerotome-derived fibroblast subtypes identified by scRNA-seq appeared to be terminally differentiated (Fig. 1G). To test whether they can contribute to new tenocytes after injury, we performed single-cell clonal analysis of three specialized fibroblast subtypes: tenocytes, notochord-associated fibroblasts, and fin mesenchymal cells. First, we asked whether unablated tenocytes remaining after laser ablation could contribute to new tenocytes along the injury site. In uninjured embryos, tenocytes did not proliferate or contribute to new tenocytes (Figs. 5C,D,F). Unlike perivascular and interstitial fibroblasts, most tenocytes (86%, 19/22) showed no change in behavior even following injury (Figs. 5B,C). Accordingly, we did not find a significant increase in clone size of traced tenocytes in injured embryos compared to uninjured controls (Fig. 5F). These results show that unablated tenocytes are not activated upon tendon injury. Like tenocytes, neither fin mesenchymal cells nor notochord-associated fibroblasts showed noticeable activation after injury (Fig. 6), with 100% of fin mesenchymal cells (7/7) and 84% of notochord-associated fibroblasts (16/19) remaining at their original location after injury (Figs. 6A-C). Furthermore, the composition and size of their clonal population did not change substantially following injury (Figs. 6D-F).

**Fig. 6.**
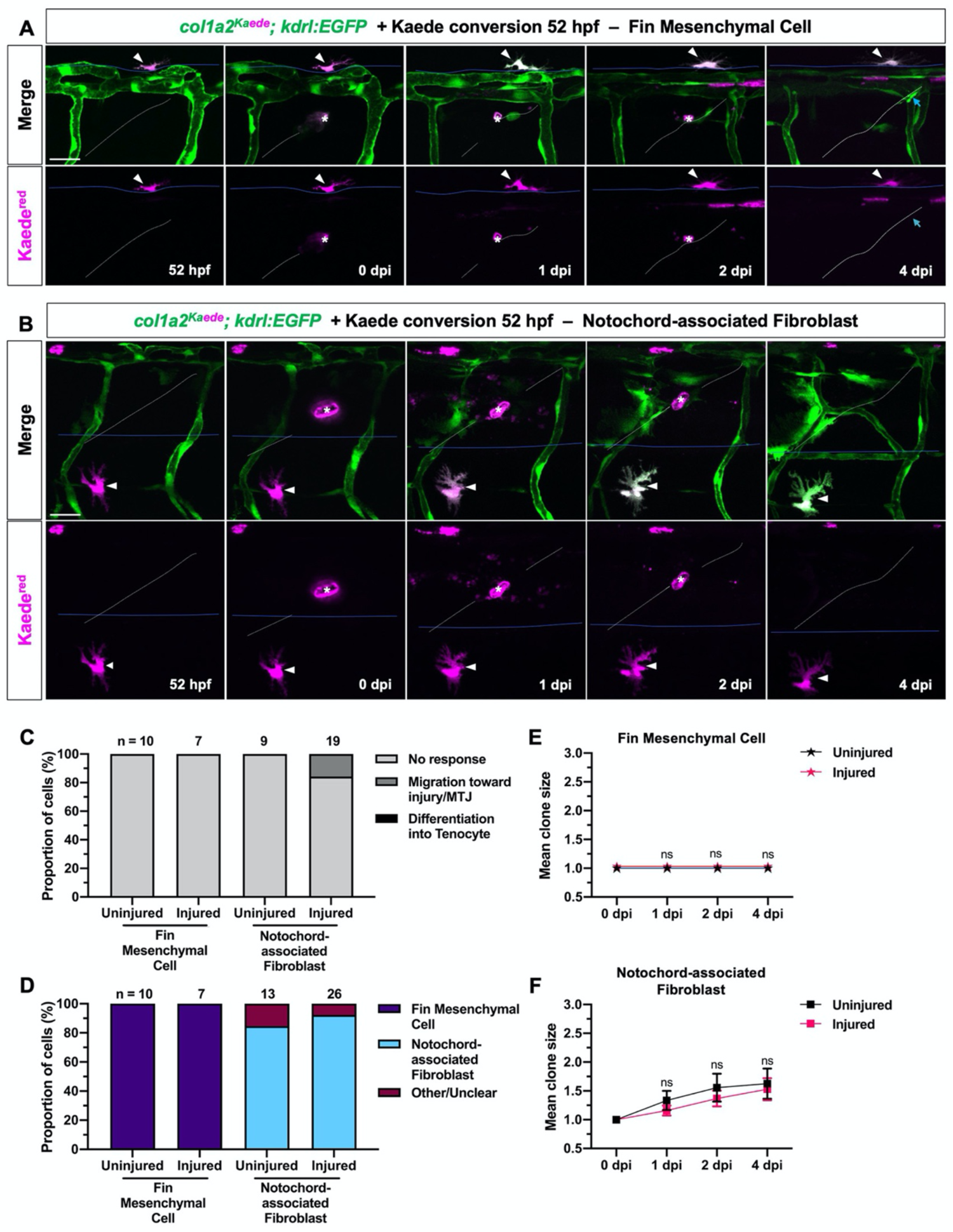
Fin mesenchymal cells and notochord-associated fibroblasts do not respond to tendon injury. (**A,B**) Clonal analysis of a single fin mesenchymal cell (A) and notochord-associated fibroblast (B) from 52 hpf to 4 dpi, as described in Fig. 4A. Both fibroblast subtypes (magenta, arrowheads) are retained within their respective anatomical locations after tendon injury. Blue lines indicate fin fold boundary in A and dorsal edge of the notochord in B. The regenerating MTJ and injury site are indicated by white dotted lines and asterisks, respectively. Blue arrows in A indicate a newly regenerated Kaede^red^-negative tenocyte. (**C-F**) Quantification of fibroblast response (C), clonal composition (D), and clone size for fin mesenchymal cells (E) and notochord associated fibroblasts (F) in injured versus uninjured embryos. Neither cell type showed notable activation after tendon injury. *n* = 10 (uninjured) and 7 (injured) fin mesenchymal cells; 9 (uninjured) and 19 (injured) notochord-associated fibroblasts. Data in E,F shown as mean ± SEM. Statistics: Sidak’s multiple comparisons (E,F). Significance: p-value >0.05 (ns). Scale bars: 25 μm.

Altogether, our single-cell clonal analysis confirms that specialized sclerotome-derived fibroblast subtypes, including tenocytes, fin mesenchymal cells, and notochord-associated fibroblasts, appear fate-committed and do not contribute to tendon regeneration.

### Inhibition of Hedgehog signaling impairs tenocyte regeneration

Our clonal analysis reveals a key role for perivascular/interstitial fibroblasts in tendon regeneration. This observation raises the question: Which cell signaling pathways drive the recruitment and differentiation of these responding fibroblasts? Interestingly, *ptc2*, a target gene of the Hedgehog (Hh) signaling pathway, was upregulated near the injury site in 100% of embryos (10/10) at 1 dpi and 64% of embryos (18/28) at 2 dpi (Fig. 7A). Consistently, 63% (17/27) and 64% (16/25) of *ptc2:Kaede; col1a2^NTR-mCherry^* embryos possessed Kaede^+^ fibroblasts near the injury site at 1 and 2 dpi, respectively (Fig. 7B). Interestingly, of all Kaede^+^ cells at the injury site, only 48% (32/67) in 1 dpi embryos and 64% (35/55) in 2 dpi embryos were also labelled by *col1a2^NTR-mCherry^*, suggesting that fibroblasts are not the only Hh-responsive cell population after tenocyte ablation. To test whether Hh signaling is important for tenocyte regeneration, we treated embryos with cyclopamine, a potent inhibitor of Hh signaling (*39*), after tendon injury (Fig. 7C). At 1 dpi, a large gap in *prelp* staining at the injured MTJ was observed in both DMSO and cyclopamine-treated embryos, suggesting new tenocytes had not emerged in either group (Figs. 7D-F). As expected, by 2 dpi this gap was significantly reduced and the *prelp* expression was expanded in DMSO-treated controls (Figs. 7D-F). Strikingly, no such change was observed in cyclopamine-treated embryos, which showed reduced *prelp* expression area and large gaps in staining at the injury site at 2 dpi, indicating compromised tenocyte regeneration in these fish (Figs. 7D-F). Similar tenocyte regeneration defects were also reproduced with another tenocyte marker, *tnmd* (Figs. S11A-C).

**Fig. 7.**
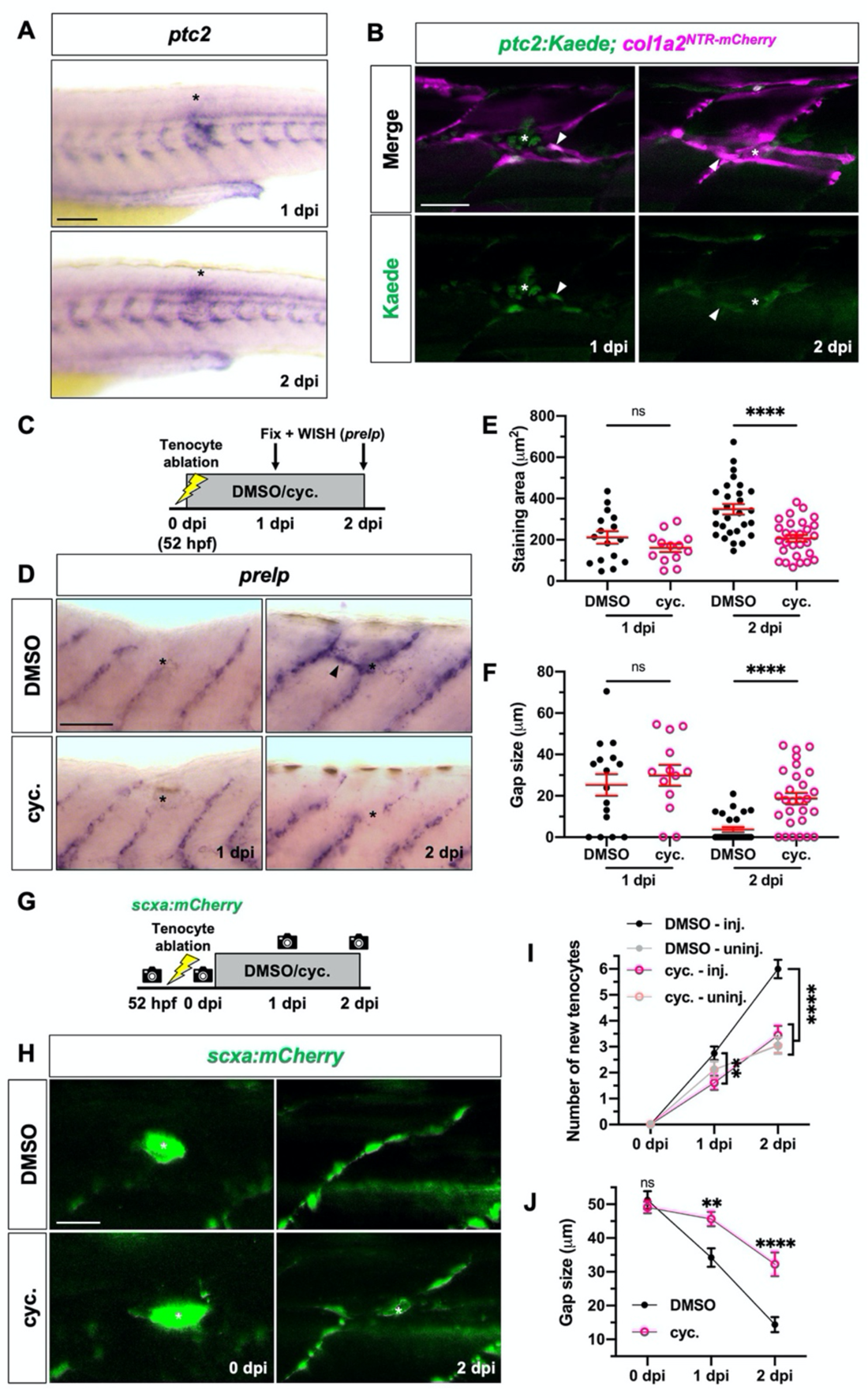
Inhibition of Hh signaling impairs tenocyte regeneration. (**A**) Whole-mount mRNA in situ hybridization showing upregulation of Hh target gene *ptc2* at the injury site (asterisks) at 1 and 2 dpi. *n* = 10/10 (1 dpi) and 18/28 (2 dpi) embryos. (**B**) Images of *ptc2:Kaede; col1a2^NTR-mCherry^*embryos at 1 and 2 dpi showing Kaede^+^mCherry^+^ fibroblasts (arrowheads) near the injury site (asterisks). *n* = 17/27 (1 dpi) and 16/25 (2 dpi) embryos. (**C**) Experimental protocol for drug treatment and staining of injured embryos. Injured embryos were incubated in DMSO or cyclopamine (cyc.) prior to fixation and in situ hybridization for *prelp* at appropriate stages. (**D**) Tenocyte marker *prelp* expression in DMSO and cyclopamine-treated embryos at 1 and 2 dpi. Ectopic *prelp* expression (arrowhead) outside the MTJ can be seen in DMSO-treated but not cyclopamine-treated fish at 2 dpi. (**E,F**) Quantification of staining area (E) and gap size (F) for drug treated embryos as described in Fig. 2A. *n* = 16 (1 dpi) and 28 (2 dpi) DMSO-treated embryos; 13 (1 dpi) and 28 (2dpi) cyclopamine-treated embryos. (**G**) Drug treatment protocol for live *scxa:mCherry* embryos in H-J. (**H**) Representative images of injured MTJs from DMSO and cyclopamine-treated embryos at 0 and 2 dpi. A notable gap between tenocytes (green) can be seen in cyclopamine but not DMSO-treated embryos. (**I,J**) Number of new tenocytes generated at injured and uninjured MTJs (I), and mean gap between tenocytes at the injury site (J) from 0-2 dpi in DMSO and cyclopamine-treated embryos. *n* = 32 (DMSO) and 28 (cyclopamine) embryos. All data are plotted as mean ± SEM. Statistics: Mann-Whitney *U* test (E,F); Sidak’s multiple comparisons (I,J). Significance: p-value >0.05 (ns), <0.01 (**), <0.0001 (****). Asterisks denote injury site in all images. Scale bars: 100 μm (A); 50 μm (B,D); 25 μm (H).

To directly visualize the effect of Hh inhibition on tenocytes, we tracked tenocyte number in live *scxa:mCherry* embryos treated with DMSO or cyclopamine from 0 to 2 dpi (Fig. 7G). Adjacent uninjured MTJs in injured fish were scored as controls. Both DMSO and cyclopamine-treated embryos showed a similar rate of tenocyte production along uninjured MTJs during the examined period, indicating that cyclopamine treatment does not impair normal tenocyte development (Fig. 7I). As expected, significantly more tenocytes were produced along injured MTJs in DMSO-treated embryos compared to uninjured MTJs in the same fish (Figs. 7H,I). By contrast, in cyclopamine-treated embryos, the rate of tenocyte production at injured MTJs was similar to uninjured MTJs in the same fish, and significantly lower than those at injured MTJs of DMSO-treated siblings (Fig. 7I). Consistently, the distance between adjacent tenocytes at the injury site was also significantly larger in cyclopamine-treated versus DMSO-treated fish (Figs. 7H,J). To confirm that these tenocyte regeneration defects were indeed due to disruption of Hh signaling, we tested two additional small molecule inhibitors of Smoothened, LDE225 (Sonidegib) (*40*) and BMS-833923 (Fig. S11D) in *scxa:mCherry* embryos. Like cyclopamine, these drugs have been shown to robustly inhibit Hh signaling in zebrafish (*41*, *42*). For each drug tested, treatment from 0 to 2 dpi resulted in reduced tenocyte production and increased gap between tenocytes at injured MTJs relative to DMSO-treated controls (Figs. S11E-G). Collectively, these results demonstrate that inhibition of Hh signaling at 0-2 dpi substantially impairs tenocyte regeneration after injury.

### Hedgehog signaling is required for the proliferation of activated fibroblasts

Next, we aimed to elucidate the mechanism by which Hh signaling mediates tenocyte regeneration. To visualize how Hh inhibition affects fibroblast behavior, we performed single-cell clonal analysis of perivascular fibroblasts in *col1a2^Kaede^; kdrl:EGFP* embryos treated with cyclopamine from 0 to 2 dpi, as previously described (Fig. 8A). As expected, most perivascular fibroblasts (91%, 10/11) were retained along the ISV in uninjured embryos treated with cyclopamine (Figs. 8B,C). After injury, 53% of perivascular fibroblasts (8/15) in DMSO-treated embryos and 67% in cyclopamine-treated embryos (18/27) migrated to the injury site between 0-2 dpi (Figs. 8B,C). However, only 7% of traced cells (2/27) in cyclopamine-treated embryos appeared to differentiate into tenocytes compared to 13% in DMSO-treated controls (2/15) (Fig. 8C). Interestingly, mean clone size at 2 dpi was significantly reduced in injured cyclopamine versus DMSO-treated embryos, suggesting reduced fibroblast proliferation (Fig. 8D). To confirm this proliferation defect, we examined EdU incorporation from 0 to 2 dpi in DMSO or cyclopamine-treated *col1a2^Kaede^*embryos (Fig. 8E). Consistent with our previous results, we observed an overall reduction in the number of EdU^+^ fibroblasts (Kaede^+^) and total EdU^+^ cells in the injury region in cyclopamine-treated embryos compared to DMSO-treated controls at 2 dpi (Figs. 8F-H). Based on these results, we conclude that although not required for the initial recruitment of fibroblasts to the injury site, Hh signaling promotes the proliferation of injury-responsive fibroblasts.

**Fig. 8.**
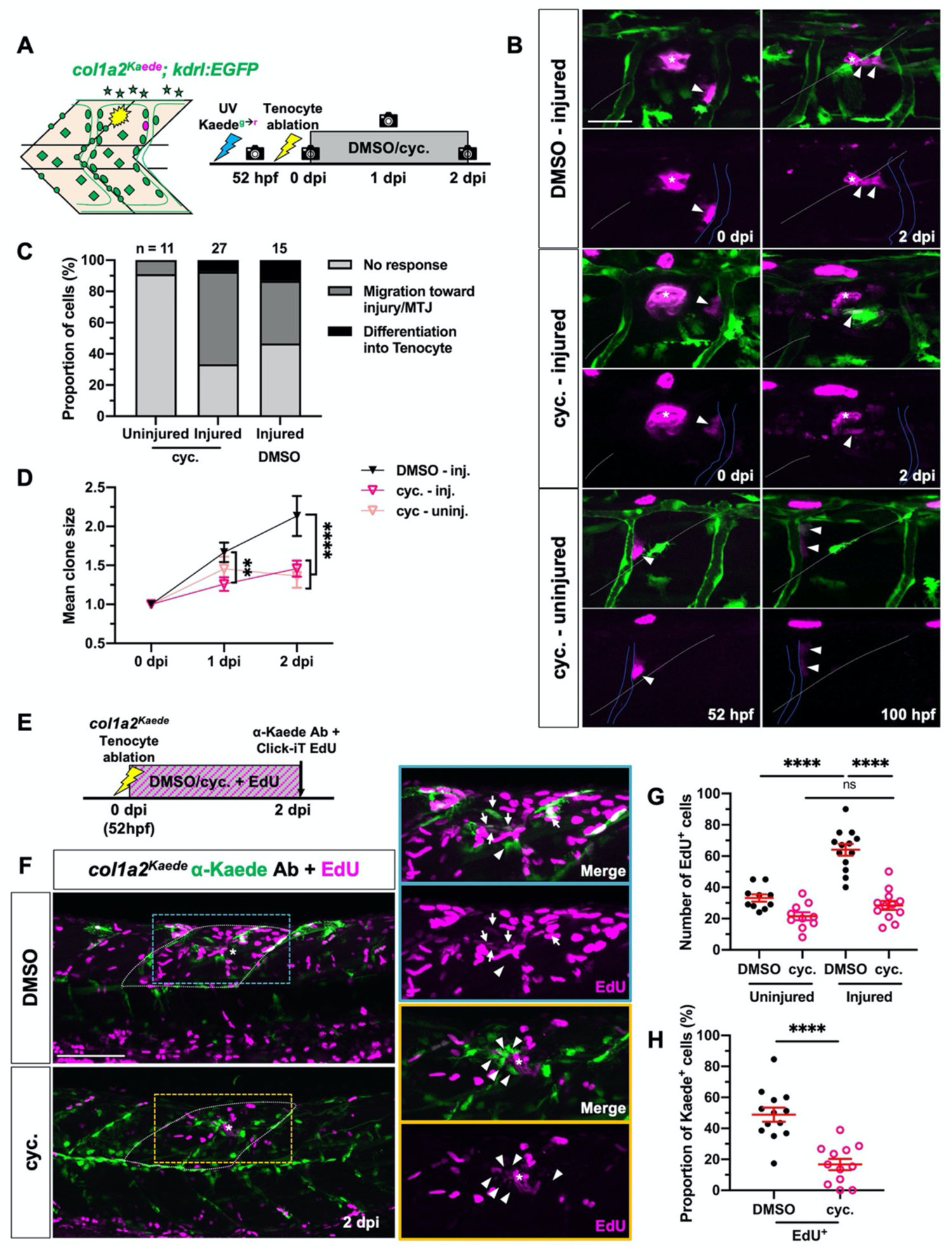
Hh signaling regulates fibroblast proliferation during tenocyte regeneration. (**A**) Experimental protocol for single-cell clonal analysis of perivascular fibroblasts in DMSO or cyclopamine-treated embryos. (**B**) Representative images of photoconverted perivascular fibroblasts and their progeny in injured DMSO and cyclopamine-treated embryos at 0 and 2 dpi, and uninjured cyclopamine-treated embryos at equivalent stages (52 and 100 hpf). Labelled clones (arrowheads) can be seen migrating from the ISVs (blue lines) to the injury site (asterisks) in both injury conditions, but are retained along the ISVs in uninjured cyclopamine-treated fish. White dotted lines indicate the MTJs. (**C,D**) Overall perivascular fibroblast response (C) and clone size (D) in DMSO or cyclopamine-treated embryos after tenocyte ablation. *n* = 15 (injured) DMSO-treated embryos; 11 (uninjured) and 27 (injured) cyclopamine-treated embryos. (**E**) Timeline for concurrent EdU incorporation and drug-treatment post injury. (**F**) Representative images from control and drug-treated embryos from E. Most fibroblasts (labelled by *col1a2^Kaede^*) surrounding the injury site (asterisks) in DMSO-treated embryos at 2 dpi were EdU^+^ (arrows), whereas majority of injury-responsive fibroblasts in cyclopamine-treated embryos showed no EdU labelling (arrowheads). EdU-labelled cells in the dorsal half of two somites immediately surrounding the injury site (white dotted lines) were counted as shown in G,H. (**G,H**) Quantification of total EdU^+^ cells (G) and proportion of EdU^+^ fibroblasts (H) at injured and uninjured regions in DMSO and cyclopamine-treated embryos. *n* = 10 (uninjured) and 13 (injured) MTJs from 13 DMSO-treated embryos; 10 (uninjured) and 12 (injured) MTJs from 12 cyclopamine-treated embryos. All data are plotted as mean ± SEM. Statistics: Mann-Whitney *U* test (G,H); Sidak’s multiple comparisons (D). Significance: p-value >0.05 (ns), <0.01 (**), <0.0001 (****). Asterisks denote injury site in all images. Scale bars: 100 μm (F); 25 μm (B).

Since Hh signaling was not necessary for fibroblast recruitment, we hypothesized that shorter duration of Hh inhibition from 1-2 dpi, after most responding fibroblasts have migrated, would be sufficient to produce tenocyte regeneration defects. To test this, we examined *prelp* expression in injured embryos treated with DMSO or cyclopamine from 0-2 dpi or 1-2 dpi (Fig. S12A). Both cyclopamine-treated groups showed comparable large gaps in staining at the injury site at 2 dpi as well as lower overall *prelp* expression area compared to DMSO-treated controls (Figs. S12B-C). Therefore, we determine 1-2 days post injury as the critical period for Hh signaling-mediated tenocyte regeneration.

## DISCUSSION

Our work combines scRNA-seq, lineage tracing, and clonal analysis to investigate fibroblast behavior and plasticity during tenocyte injury repair in zebrafish (Fig. 9). Our experiments provide three key findings. First, we show that the sclerotome gives rise to six transcriptionally distinct fibroblast subtypes, including perivascular/interstitial fibroblasts, tenocytes, fin mesenchymal cells, notochord-associated fibroblasts, stromal reticular cells and DLAV fibroblasts. Second, using a tenocyte regeneration model, we show that perivascular and interstitial fibroblasts, but not other fibroblast subtypes, become activated and contribute to new tenocytes after injury. Finally, we establish a timeline of axial tenocyte regeneration in zebrafish and demonstrate a critical role for the Hh signaling pathway in regulating fibroblast proliferation during tendon injury repair. Altogether, our work highlights the differences in plasticity amongst fibroblast populations in tissue repair and provides key insights into the molecular and cellular basis of tendon regeneration.

**Fig. 9.**
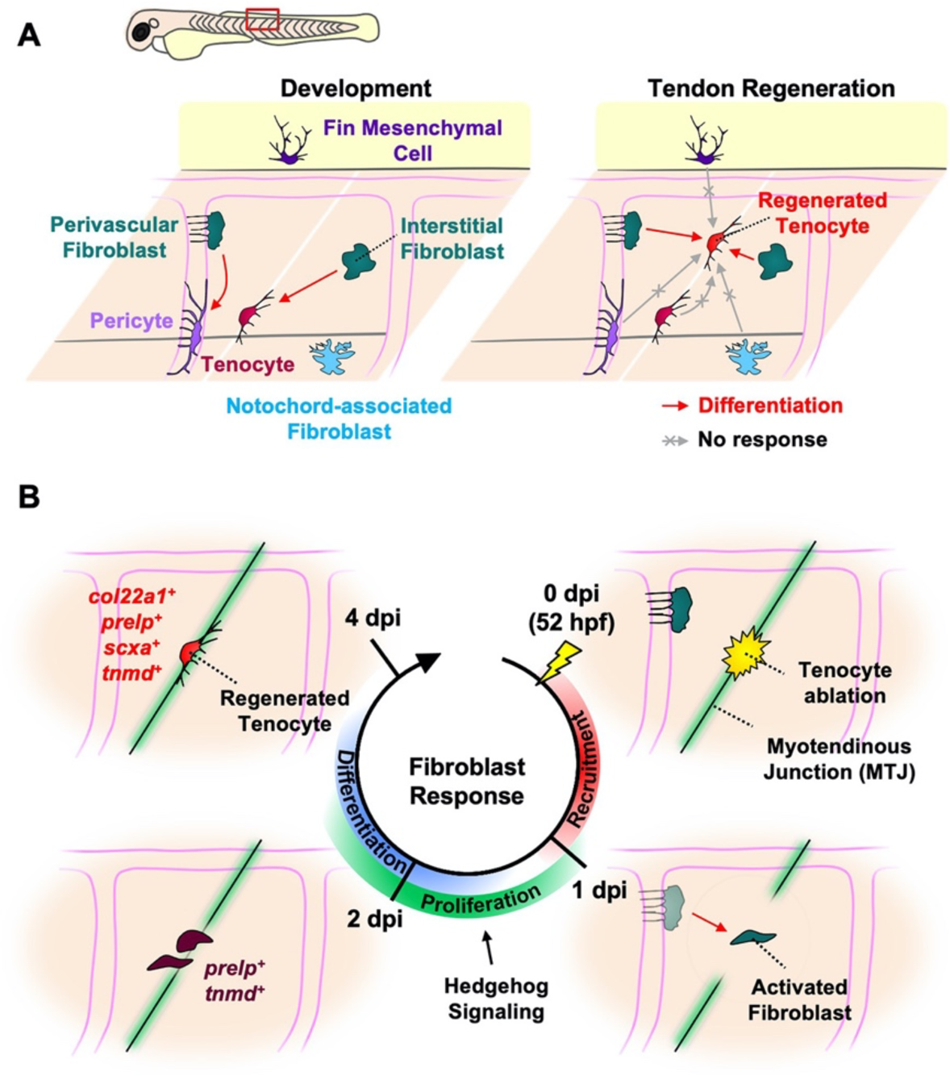
Model of fibroblast plasticity and tendon regeneration in zebrafish. (**A**) Schematic showing distinct plasticity of different sclerotome-derived fibroblast populations in the zebrafish trunk at 52 hpf, including progenitor-like perivascular/interstitial fibroblasts, as well as specialized tenocytes, fin mesenchymal cells, and notochord-associated fibroblasts. During normal development, perivascular fibroblasts act as pericyte precursors, while interstitial fibroblasts function as tenocyte progenitors. By contrast, both perivascular and interstitial fibroblasts contribute to tenocyte regeneration upon tendon injury, while perivascular fibroblast-derived pericytes as well as specialized fibroblasts do not exhibit such regenerative plasticity. (**B**) Timeline of fibroblast response to tenocyte ablation. After tenocyte ablation, perivascular fibroblast-mediated tenocyte regeneration proceeds in three partially overlapping phases. First, perivascular fibroblasts are recruited to the injured myotendinous junction from the vasculature between 0 and 1 dpi. Second, the activated fibroblasts then undergo 1-2 rounds of cell division in a Hedgehog signaling-dependent manner starting at 1 dpi, and concurrently upregulate tenocyte marker genes, *prelp* and *tnmd*. Finally, by 4 dpi, a subset of these differentiating fibroblasts matures into new tenocytes extending characteristic cellular processes into the newly regenerated MTJ.

### Different fibroblast subtypes show distinct plasticity during tissue regeneration

Because of their widespread distribution and active role in tissue maintenance and repair, it has often been hypothesized that fibroblasts can function as tissue-resident progenitors for specialized cell types. This hypothesis has been supported by numerous *in vitro* studies in which dermal and iPSC-derived fibroblasts have been successfully differentiated into mesenchymal cell types including chondrocytes, adipocytes, and osteoblasts (*43–45*). In fact, the remarkable similarity of cultured fibroblasts to mesenchymal stem cells in expression profile and differentiation potential has led to fibroblasts often being termed ‘mesenchymal progenitors’ and ‘stromal cells’ (*46*, *47*). Although *in vivo* evidence of fibroblast plasticity is limited, certain myeloid-derived fibroblasts and tissue-resident *Hic1^+^*fibroblasts have been shown to transdifferentiate into adipocytes and specialized skin fibroblasts after skin injury in mice (*48–50*). Similarly, *Hic1^+^* mesenchymal progenitors have also been shown to contribute to new tenocytes but not muscle fibers, following muscle injury in adult mice (*51*). More recently, single-cell profiling studies have also identified ‘stem-like’ fibroblasts in the adult mouse and human tissues using computational lineage inference techniques (*9*, *13*). These findings raise the question whether ‘progenitor-like’ fibroblasts are a general phenomenon extending to other tissue types and organisms.

Based on our results, this indeed appears to be the case in zebrafish. Using single-cell clonal analysis, we demonstrate that interstitial fibroblasts give rise to tenocytes during development, while both perivascular and interstitial fibroblasts do so after tendon injury (Fig. 9A). These perivascular/interstitial fibroblasts in zebrafish show remarkable similarities to stem-like ‘universal’ fibroblasts identified in adult mice (*13*). First, like universal fibroblasts, perivascular and interstitial fibroblast are more widely distributed compared to other fibroblast subtypes in the zebrafish trunk. Second, perivascular fibroblasts are associated with intersegmental vessels, similar to *Dpt^+^* universal fibroblasts which are enriched near the vasculature in the lung and small intestine (*13*). This suggests that perhaps blood vessels provide a unique niche for stem-like fibroblasts. Third, *Dpt^+^* universal fibroblasts contribute to ‘activated’ myofibroblasts in response to wounding by subcutaneous tumor implantation (*13*). Similarly, perivascular/interstitial fibroblasts show increased migration and proliferation after injury, characteristic of activated myofibroblasts. Finally, universal fibroblasts are predicted to be precursors for tissue-specific ‘specialized’ fibroblasts (*13*). We demonstrate that perivascular/interstitial fibroblasts give rise to at least one such population (tenocytes) *in vivo,* although determining whether they can contribute to other types of fibroblasts requires further experimentation. Based on these observations, we propose that perivascular and interstitial fibroblasts represent unique ‘progenitor-like’ fibroblasts in zebrafish. Whether similar progenitor-like fibroblasts are present in other zebrafish tissues and whether their plasticity is maintained into adulthood are exciting questions for future studies.

### Perivascular fibroblasts but not pericytes respond to tendon injury in zebrafish

The vertebrate vasculature is supported by a network of mural cells, comprising vascular smooth muscle cells and pericytes, and blood vessel-associated fibroblasts (*52–57*). A subset of these perivascular cells has been shown to become activated and deposit fibrotic ECM in response to injury to the brain (*58–61*), spinal cord (*61–64*), nerve (*65*), muscle (*66*), and major organs (*67*) in mice. However, cellular heterogeneity and the lack of specific markers have made the precise identification of injury-responsive perivascular populations challenging. Owing to their location along minor blood vessels and expression of pericyte marker *pdgfrb*, a few studies have identified these injury-responsive cells in the mouse central nervous system (CNS) as ‘type A’ pericytes (*58*, *61*, *62*). However, pericyte markers including *pdgfrb* are not specific, and blood vessel-associated fibroblasts have previously been found to express low levels of *pdgfrb* (*27*, *52*, *55*, *60*, *68*, *69*). Furthermore, responding perivascular cells in several mouse CNS injury models are labelled by *Col1a1* and *Pdgfra*, classic fibroblast markers not typically expressed in pericytes (*59*, *63*, *66*). Interestingly, we find both markers are also expressed in perivascular fibroblasts identified in our study. The question of regenerative perivascular cell identity is further complicated by conflicting accounts of mural cell plasticity. While pericytes have been reported to function as progenitors for numerous mesenchymal lineages like chondrocytes, adipocytes, and osteoblasts *in vitro* (*67*, *70*), they show no such differentiation potential *in vivo* (*71*). Therefore, despite extensive research, the true regenerative potential and plasticity of blood vessel-associated populations is still unclear.

Perivascular fibroblasts and pericytes comprise the major blood vessel-associated cells surrounding intersegmental vessels in the zebrafish trunk (*27*, *72*). Despite their shared location, we have previously shown that these perivascular fibroblasts and pericytes are distinct cell types with unique morphology and gene expression patterns (*27*). In fact, we find that perivascular fibroblasts function as precursors for ISV pericytes during vascular development (*27*). In this study, we utilize live imaging and lineage tracing to show that perivascular fibroblasts can also give rise to new tenocytes upon tendon injury. Pericytes, despite being derived from perivascular fibroblasts, show no such regenerative or differentiation potential upon tenocyte ablation.

Together, these findings clarify that perivascular fibroblasts but not pericytes represent progenitor-like cells associated with blood vessels in the zebrafish trunk. Our conclusions are also supported by a recent study in adult zebrafish, where *col1a2^+^* fibroblast-like perivascular cells, but not pericytes, contribute to new mural cells along regenerating vasculature following fin amputation (*73*). Yet another study shows that spinal cord injury in larval zebrafish triggers the recruitment of ISV-associated *pdgfrb^+^* cells to the lesion site, where they promote axon regeneration (*74*). Since *pdgfrb* is expressed in perivascular fibroblasts (*27*), it is conceivable that injury-activated cells in this model also represent perivascular fibroblasts rather than pericytes.

### Zebrafish is an excellent model to study tendon regeneration

Tendons are vital connective tissues connecting muscle to bone and tendon injury is often associated with devastating physical disability and reduced mobility (*75*). Unlike mammals, we and others find that zebrafish can completely regenerate their tendon structure after injury (*76–78*). Using this powerful zebrafish model, we identify sclerotome-derived perivascular/interstitial fibroblasts as the cellular source of newly regenerated tenocytes upon tendon injury. Next, we demonstrate that fibroblast response to tendon injury is separated into three overlapping phases (Fig. 9B): recruitment at 0-1 dpi, proliferation at 1-2 dpi, and differentiation at 2-4 dpi. Finally, we demonstrate the proliferation of activated fibroblasts is mediated by the Hh signaling pathway. Altogether, our work provides important insights into fibroblast behavior during tendon regeneration and provides new therapeutic targets to facilitate tissue repair.

Several injury-responsive tendon stem/progenitor populations have been identified in mice, although their response has been found to vary with age, injury mechanism, and tendon type (*79*, *80*). For instance, uninjured *Scx^+^* tenocytes have been shown to modulate wound healing after transection of the flexor digitorum longus (FDL) tendon in adults (*81*, *82*) and the Achilles tendon in neonates but not in adults (*83*). Similarly, other studies have shown *αSMA^+^* (*84*), *Tppp3^+^Pdgfra^+^* (*85*), and *Bglap^+^*(*86*) populations within the tendon sheath contribute to injury repair in several types of tendons in adult mice. However, because these studies examine different tendons and utilize distinct markers, the level of overlap between these progenitor populations remains unclear. Much less is known regarding tendon regeneration in zebrafish. One recent study shows that in larval zebrafish following the genetic ablation of tenocytes, new tenocytes in the sternohyoideus (SH) tendon arise from neighboring progenitors derived from the neural crest (*sox10^+^*) and mesodermal (*nkx2.5^+^*) lineages (*77*). Interestingly, residual *scxa*^+^ tenocytes are also found to contribute to new cells in the SH tendon upon partial tenocyte ablation (*77*). Consistently, uninjured tenocytes have been shown to become activated and give rise to new tenocytes after maxillary superficial tendon transection in adult zebrafish jaw (*76*). Similarly, lineage tracing shows that mature *scxa^+^*ligamentocytes contribute to the complete regeneration of jaw joint ligament following transection in adult zebrafish (*78*). In contrast to these studies, we find that unablated tenocytes show no active response to laser-mediated tendon injury. Instead, new tenocytes are derived from preexisting perivascular/interstitial fibroblasts in our model. Three scenarios might explain these discrepancies. First, tendon regeneration mechanism in zebrafish could vary at different developmental stages (embryos versus larvae/adults). Perivascular/interstitial fibroblasts may lose their regenerative plasticity past the embryonic stages, necessitating the involvement of mature tenocytes in tendon repair. Second, the different regenerative mechanisms observed might reflect the distinct developmental origins of tenocytes in zebrafish jaw and trunk tendons: cranial tenocytes are neural crest-derived (*22*), while axial tenocytes are sclerotome-derived (*24*). Lastly, different types of tendon injury (highly localized laser ablation versus animal-wide tenocyte ablation or tendon transection) might activate distinct progenitors.

Several signaling pathways have also been implicated in different tendon repair models, including TGFβ (*86*, *87*), PDGFα (*85*), Wnt (*88*) and Shh (*86*) signaling in mice, and BMP signaling in zebrafish (*77*). In our model, we find that Hh signaling is induced in some activated fibroblasts at the injury site after tenocyte ablation.

Consistently, inhibition of Hh signaling severely hampers the generation of new tenocytes at injured MTJs but does not affect normal tenocyte development and maintenance along uninjured junctions. Finally, we demonstrate these tenocyte regeneration defects result from compromised proliferation of activated fibroblasts. Interestingly, Wang et al. describe a similar role for Hh signaling in Tibialis anterior and Achilles tendon injury repair in mice (*86*). Consistent with our observations, they report upregulation of Hh pathway components, *Shh*, *Gli1*, and *Ptch1*, within injured tendon tissue (*86*). Furthermore, conditional deletion of the Hh pathway component *Smo* in tenocyte progenitors leads to reduced progenitor proliferation and impaired tendon regeneration, while constitutive activation of Hh signaling has the opposite effect (*86*). However, what is less clear is the source of the Hh ligand after tendon injury. The notochord and floor plate, which normally express Hh genes during zebrafish development, might be the source for Hh ligands following tenocyte ablation as well. Interestingly, Romero et al. demonstrate that damaged-induced reactive oxygen species (ROS) after tail excision in 72 hpf zebrafish trigger the expression of *shha* and *ihhb* in notochord cells at the injury site, which is sufficient to induce Hh signaling to promote tail regeneration (*41*). Similarly, H_2_O_2_, the primary ROS produced following injury, is shown to induce Hh signaling in a reciprocal manner during caudal fin regeneration in adult zebrafish (*89*). Injury-induced ROS production might therefore be a possible trigger for Hh signaling in tenocyte regeneration.

In summary, our work demonstrates that different fibroblast subtypes exhibit distinct regenerative plasticity during tendon injury repair. Progenitor-like fibroblasts might represent an evolutionarily conserved population that can be harnessed for regenerative medicine.

## MATERIALS AND METHODS

### Zebrafish strains

All animal experiments were conducted in accordance with the principles outlined in the current Guidelines of the Canadian Council on Animal Care. All protocols were approved by the Animal Care Committee of the University of Calgary (#AC17-0128 and #AC21-0102). Fish older than 24 hpf were grown in fish water containing 1-phenyl 2-thiourea (PTU) to prevent pigmentation. The following transgenic lines were utilized in our study: *TgBAC(col1a2:Gal4-VP16)ca102* (*24*), *Tg(kdrl:EGFP)la116* (*90*), *TgBAC(nkx3-1:Gal4-VP16)ca101* (*24*), *TgBAC(pdgfrb:Gal4FF)ca42* (*38*), *TgBAC(ptc2:Kaede)a4596* (*91*), *Tg(scxa:mCherry)K301* (*35*), *Tg(UAS:Cre-ERT2)ca105* (*37*), *Tg(UAS:Kaede)s1999t* (*92*), *Tg(UAS:NTR-mCherry)c264* (*92*), and *Tg*(*-3.5ubb:LOXP-GFP-LOXP-mCherry)cz1701* (*Ubi:Switch*) (*93*). The mosaic *col1a2:Gal4; UAS:Kaede* line was maintained by raising embryos with more mosaic Kaede expression.

### Tissue dissociation and cell sorting

About 140 *nkx3-1^NTR-mCherry^* embryos at 52 hpf were used for single-cell RNA sequencing. Embryos were anesthetized in fish water containing 0.4% tricaine and maintained in an ice bath during dissection. Embryos were dissected at the beginning of the yolk extension using a surgical scalpel and trunk regions were collected for dissociation. To prepare the single-cell suspension, trunk tissues were incubated in 0.25% Trypsin and 1 mM EDTA solution at 28.5°C for 20 minutes with gentle shaking at 300 rpm. Dissociation was stopped by adding 100 μl FBS (Fetal Bovine Serum) and 1.25 μl 0.8M CaCl_2_. Dissolved tissue was subsequently washed 3 times with 1% FBS solution in 1x DPBS (Dulbecco’s phosphate-buffered saline, ThermoFisher Scientific) and passed through 75 μm and 35 μm cell strainers to remove debris.

Approximately 40,000 live, mCherry^+^ cells were collected by fluorescence-activated cell sorting (FACS) using the BD FACSAria. Gating was set to exclude doublets and debris, and dead cells (Invitrogen Fixable Viability Dye eFluor 780) prior to sorting for mCherry as shown in Fig. S1A. Sorted cells were centrifuged at 500g for 8 minutes and resuspended in 50 μl HBSS (Hank’s Balanced Salt Solution) with 2% BSA (Bovine Serum Albumin). Cell quality and number were verified visually.

### Cell separation, library preparation and sequencing

Approximately 15,000 live, mCherry^+^ cells were loaded into the 10X Genomics Chromium Controller and processed using the 10X Genomics 3’ gene expression profiling droplet-based scRNA-seq technology (v3.1 chemistry). All steps were performed according to manufacturer’s protocol. cDNA was amplified for 12 cycles and index primer addition PCR used 14 cycles. Sequencing was performed using the Illumina NovaSeq S2 flow cell using at the Centre for Health Genomics and Informatics (CHGI) at the University of Calgary.

### Computational analysis

To prepare a custom zebrafish transcriptome, a FASTA sequence for the *mCherry* gene was appended to the *D. rerio* GRCz11 reference genome and the combined genome file was indexed with V4.3.2 zebrafish transcript annotations (*94*) using CellRanger mkgtf and mkref software. FASTQs were aligned to the transcriptome using the CellRanger count pipeline with default parameters. All pipelines were executed using the 10X Genomics CellRanger v5.0.0 software (*95*). A total of 3171 cells with 90,583 mean reads per cell were detected post alignment.

The raw counts matrix was imported into R and standard Seurat v.4.0.0 protocols were used for downstream quality control, dimensionality reduction, cell clustering and marker identification (*96*). Cells were filtered prior to downstream analysis using the following quality control metrics: number of genes expressed >300 and <5000, percentage of mitochondrial genes < 10%, and log_10_(Genes/UMIs) >0.8. Additionally, genes expressed in <10 cells were excluded. Normalization of the filtered dataset was performed using the ‘SCTransform()’ function regressing for percentage of mitochondrial genes. Top 40 PCAs were used to generate the UMAP plot and a resolution of 0.8 was selected for clustering. Markers for each cluster were identified using the ‘FindAllMarkers()’ function using default parameters. Clusters of interest were then subsetted and renormalized and reanalyzed. List of ‘core matrisome’ genes for Fig. 1C was downloaded from Nauroy et al (*31*). Top markers for fibroblast subclusters were identified with the ‘FindAllMarkers()’ function with the following parameters: ‘only.pos = TRUE’, ‘logofc.threshold = 0.25’, and ‘min.diff.pct = 0.3’.

The .loom file containing Unspliced/Spliced count matrices for RNA velocity analysis was generated using the velocyto 0.17.16 command line tool (*97*). Sparse matrix, metadata and embeddings from the analyzed Seurat object were then imported into Python and combined with the .loom file to generate anndata object. RNA velocity analysis was then performed on the imported dataset using the scVelo python package using the ‘dynamical’ mode as recommended (*98*).

### In situ hybridization and immunohistochemistry

Whole-mount in situ hybridization and antibody staining were performed using standardized protocols. For double staining, fluorescent in situ hybridization was performed first followed by antibody staining. For anti-Thbs4b antibody staining, embryos were fixed directly in 95% methanol/5% glacial acetic acid. For all other staining, embryos were fixed in 4% formaldehyde in PBS (phosphate-buffered saline) for 18+ hours and then transferred to 100% methanol. The following digoxigenin (DIG) labeled probes were used in this study: *ca6*, *ccl25b*, *ccn2b*, *cdh11*, *cilp*, *clec19a*, *col22a1, crispld1b*, *flbn1*, *hapln1a*, *hgd*, *hpdb*, *nkx3-2*, *pcdh18b*, *prelp*, *ptc2*, *sfrp1a*, *tnmd*, *wnt11*, and *zic1*. The primary and secondary antibodies used in this study are listed in Table S3.

### EdU labeling

Invitrogen Click-iT™ EdU Cell Proliferation Kit (Catalog #C10338) was used for all EdU incorporation assays. For EdU pulsing experiment in Fig. S6C, embryos were incubated in 500 μM EdU in PTU fish water from 52 to 53 hpf. For longer term treatment post injury in Figs. 8E-H, embryos were incubated in 100 μM EdU in PTU water with 100 μM cyclopamine or DMSO from 0 to 2 dpi. After treatment, embryos were washed and fixed in 4% formaldehyde in PBS for 24 hours and then transferred to 100% methanol. Click-iT reaction was performed according to manufacturer’s guidelines.

### Quantification of cycling fibroblasts

To quantify proliferative fibroblast populations in Fig. S6D, PHH3 or EdU positive fibroblasts (Kaede^+^) were manually counted in each embryo. Fibroblast subtype identity was determined based on morphology and anatomical location. Counting of Kaede^+^EdU^+^ fibroblasts and total EdU^+^ cells in Figs. 8G-H was done manually using the ‘CellCounter’ plugin in ImageJ.

### Tenocyte ablation

Embryos were mounted as described in the ‘confocal imaging’ section. Using the 25x objective of the Leica TCS SP8 multiphoton microscope at 48x zoom, a 9.23 μm x 9.23 μm region at the dorsal region of a single myotendinous junction (MTJ) in each embryo was targeted for injury. Injury was induced by exposing the selected region to 900 nm laser at 75-85% laser strength for 1 frame.

### Imaging and quantification of in situ hybridization

After whole-mount in situ hybridization, embryos were mounted in 100% glycerol and imaged using the Zeiss AxioImager. Staining area was measured as depicted in Figs. 2A, S11A and S12A using Fiji software (*99*). Briefly, images were converted to 8-bit format and the dorsal section of 2 somites adjacent to the injury site was selected for quantification using the ‘polygon selections’ tool. Stained pixels were highlighted by adjusting image threshold and total area of highlighted pixels in the selected area was measured. Gap size was quantified as depicted (Figs. 2A, S11A, S12A) using the ‘line selection’ tool in Fiji.

### Confocal imaging

Confocal imaging was performed with the Olympus FV1200 confocal microscope using the 20x objective or the Leica TCS SP8 multiphoton microscope using the 25x objective. For live imaging, embryos were anesthetized in 0.4% tricaine solution and mounted in 0.8% low melting point agarose. To follow embryos from 0-4 dpi, embryos were maintained in individual wells in 24-well plates and returned to the appropriate well after imaging. Timelapse movies were generated by repeatedly imaging the region of interest every 12-20 minutes from 0-24 hours post injury (hpi). Movies were compiled using the Olympus Fluoview software and manually annotated using Fiji (*99*). All Images were processed using the Fiji software (*99*). Tenocyte number in live embryos was counted manually at each stage. Gap between adjacent tenocytes in live embryos was quantified using the ‘line selection’ tool in Fiji.

### Kaede photoconversion

Photoconversion of Kaede protein from Kaede^green^ to Kaede^red^ was performed using the 20x objective on the Olympus FV1200 confocal microscope. To convert all fibroblasts in a large area as shown in Fig. 3A, a region of 500×400 pixels spanning two somites was scanned with the 405 nm laser for 8 frames at 75-80% laser strength. For conversion of individual fibroblasts for clonal analysis in Figs. 4, 5, 6, 8, and S10, a circular region of 10 pixel diameter within the desired fibroblast was exposed to 405 nm laser for 4-6 seconds at 10% laser strength.

### Quantification of single-cell clonal analysis

Single-cell clonal populations of fibroblasts were quantified using three metrics as described in Fig. 4B. First, cell behavior was manually tracked from 0-4 dpi. Fibroblast response was further categorized into (a) migration toward injured MTJ, (b) differentiation into tenocyte, and (c) no response. Second, the pooled Kaede^red^-positive clonal population from all traced cells for each fibroblast subtype was scored for fibroblast composition at the desired stages post injury. Fibroblast subtype identity was determined based on anatomical location and morphology. Overall distribution of fibroblast subtypes across all clonal populations was graphed. Third, clone size was determined by manually counting number of Kaede^red^ cells at each stage from 0-4 dpi. Clone size was graphed as mean ± S.E.M. across all cells traced.

### Drug treatment

For Cre-mediated lineage tracing of sclerotome-derived fibroblasts, embryos were incubated in PTU water containing 10 μM 4-hydroxy tamoxifen (4-OHT) (Sigma Aldrich, H7904) from 30 to 52 hpf (Fig. 3E). After treatment, 4-OHT was washed off and treated embryos were saved for imaging and tenocyte ablation.

For inhibition of Hh signaling, embryos were incubated in 100 μM cyclopamine (Cedarlane Labs, C988400-50), 15 μM BMS-833923 (Cedarlane labs, A3258-10), or 75 μM Sonidegib NVP-LDE225 (Selleckchem, S2151) in PTU water at the appropriate stages post injury. Control embryos were incubated in equivalent concentrations of DMSO in PTU water.

### Plotting and statistical analysis

Plotting and statistical analysis for single-cell RNA sequencing data was performed using appropriate Seurat plotting functions with the ggprism extension for ggplot2 All other graphs and statistical analysis were generated using the GraphPad PRISM software. Data were plotted with mean ± S.E.M. indicated. For paired data in tenocyte tracing (Figs. 2H, 7I, S11E, S11F), gap size tracing (Figs. 7J, S11G), and clone size experiments (Figs. 4F, 4G, 5E, 5F, 6E, 6F, 8D), significance was calculated by 2-way ANOVA Sidak’s multiple comparisons test, as recommended by GraphPad PRISM. For all other data, significance was calculated by performing the non-parametric Mann-Whitney *U* test with two-tailed p values: p > 0.05 (not significant, ns), p < 0.05 (*), p < 0.01 (**), p < 0.001 (***) and p < 0.0001 (****).

## ACKNOWLEDGMENTS

We thank the zebrafish community for providing probes and reagents; Jenna Galloway for sharing the *scxa:mCherry* line; James Gagnon for discussion on scRNA-seq analysis; Sarah Childs for sharing transgenic lines and providing critical input on this project; Katrinka Kocha for fish care and assistance with embryo dissection for scRNA-seq; members of the Huang laboratory for discussions. We also thank the Center for Health Genomics and Informatics (CHGI) at the University of Calgary for providing sequencing and high-performance computing services.

## Funding

This study was supported by following grants:

Canadian Institutes of Health Research PJT-169113 (PH)

Canada Foundation for Innovation John R. Evans Leaders Fund Project 32920 (PH) Startup Fund from the Alberta Children’s Hospital Research Institute (ACHRI) (PH) Eyes High Doctoral Recruitment Scholarship from the University of Calgary (AMR)

## Author Contributions

Conceptualization: AMR, PH

Methodology: AMR, PH

Investigation: AMR, NLR, EL, PH

Visualization: AMR, PH

Resources: JB, SL, PH

Writing – Original Draft: AMR, PH

Writing – Review & Editing: AMR, NLR, EL, JB, SL, PH

Supervision: PH

Project Administration: PH

Funding Acquisition: PH

## Data and Materials Availability

All data are available in the main text or the supplementary materials. Raw FASTQs files and the processed count matrix for scRNA-seq can be accessed at the NCBI GEO database with accession number GSE229939. The original code is available at Dryad (https://doi.org/10.5061/dryad.fttdz090c) and Github (https://github.com/hpeng031/Sclerotome).

## SUPPLEMENTARY MATERIALS

**Fig. S1.**
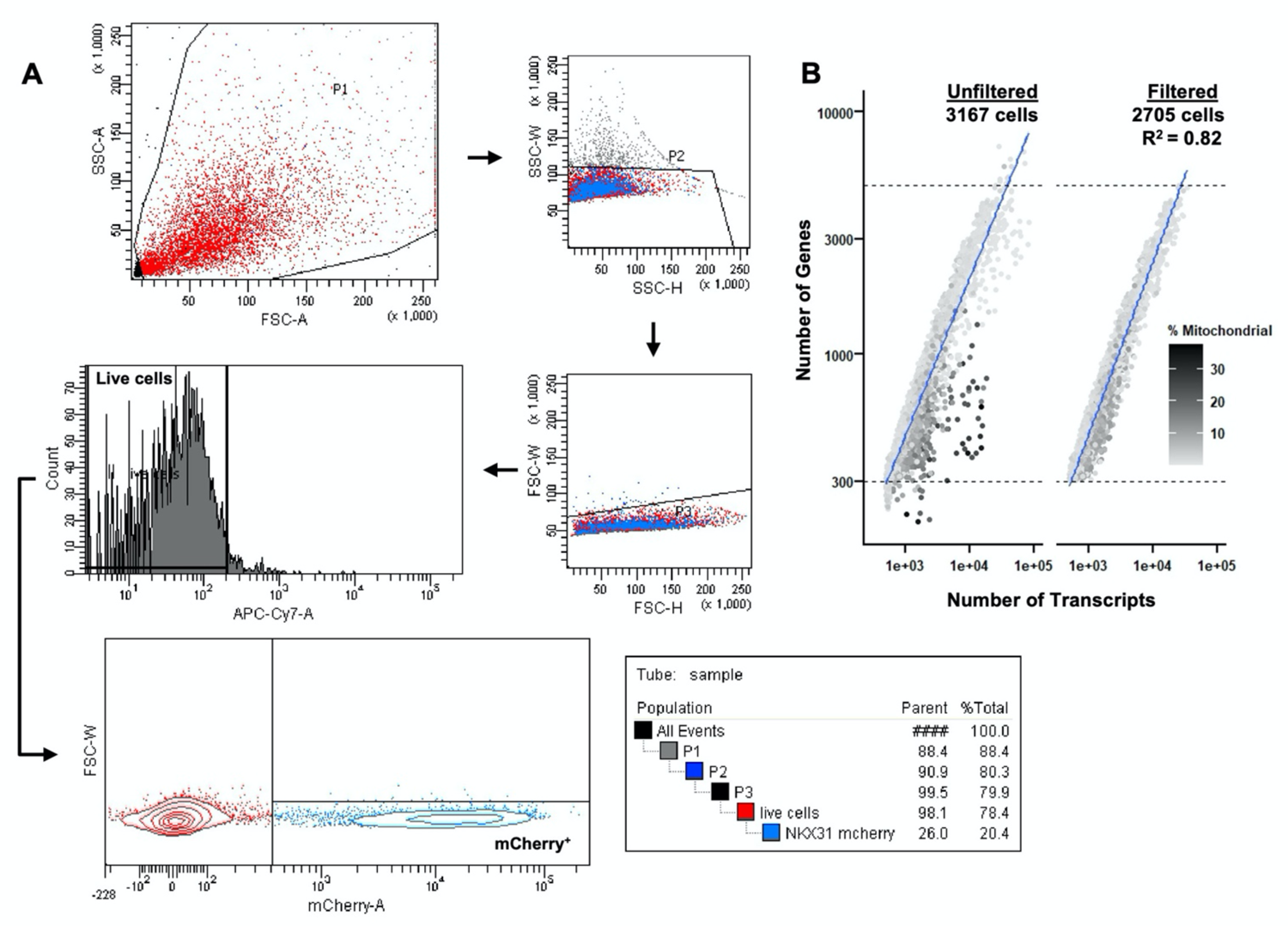
Quality control for single-cell RNA sequencing. Related to Fig. 1. (**A**) FACS gating protocol to exclude debris (P1), multiplets (P2/3) and sort live mCherry^+^ cells for input into the 10X Chromium Controller. (**B**) Total number of genes (nFeatures) versus total transcripts (nUMIs) detected per cell in the sequenced dataset before and after filtering for quality control metrics. Grading denotes concentration of mitochondrial gene transcripts per cell. A final post-filtering dataset of 2705 high quality cells was used for clustering and analysis.

**Fig. S2.**
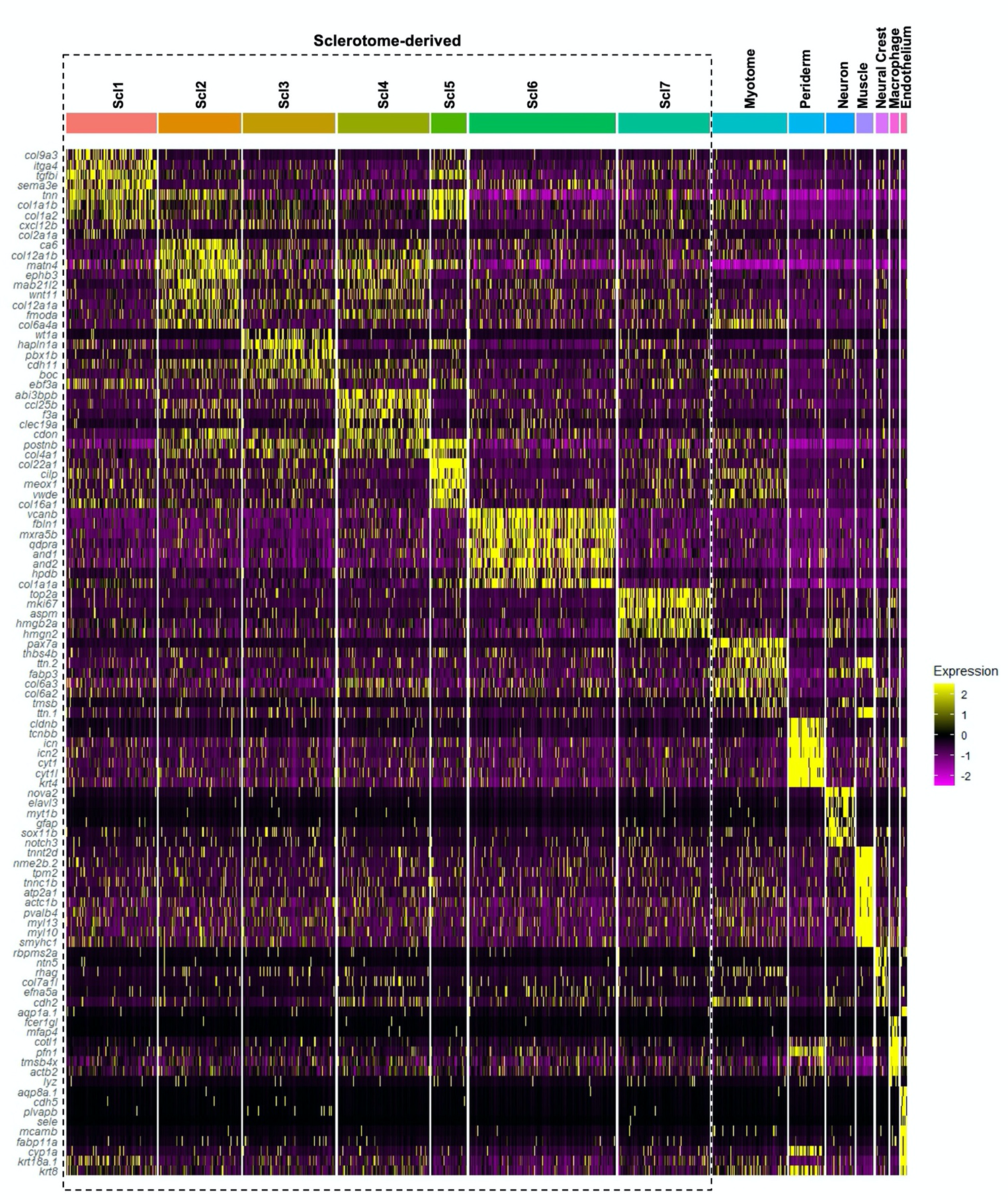
Identification of top markers for cluster annotation. Related to Fig. 1. Heatmap showing expression profile of top 7-10 markers identified for each cluster from Fig. 1B. Cluster identity was determined based on expression of previously identified marker genes.

**Fig. S3.**
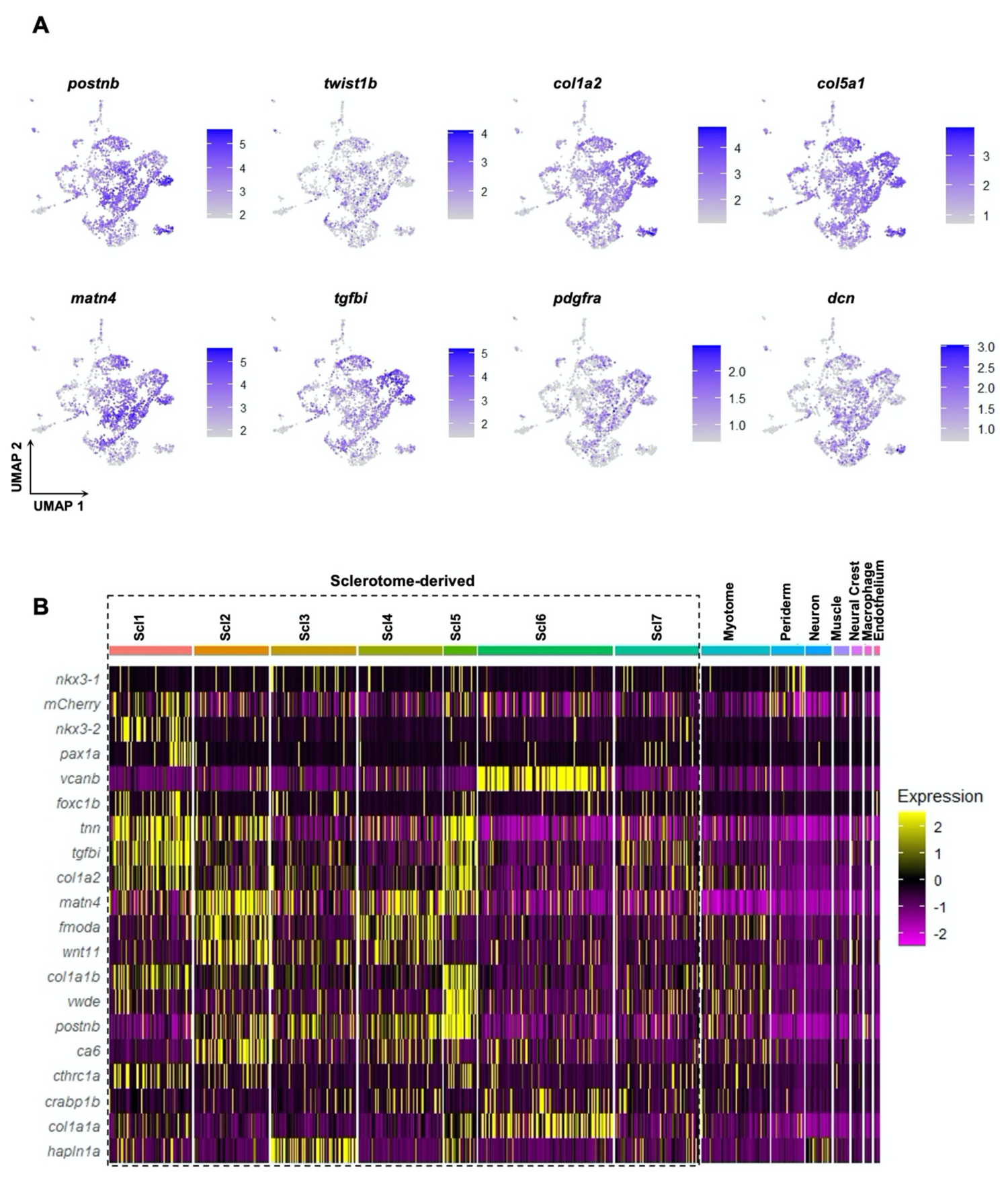
Identification of sclerotome-derived fibroblasts. Related to Fig. 1. (**A**) Feature plots showing high expression of known sclerotome (*postnb*, *twist1b*, *matn4*, *tgKi*) and pan-fibroblast (*col1a2*, *col5a1*, *pdgfra*, *dcn*) marker genes in sclerotome-derived fibroblast clusters. (**B**) Expression heatmap of previously known sclerotome markers of all clusters from Fig. 1B. Higher general expression can be seen in cells from clusters Scl1-7 representing sclerotome-derived fibroblasts (dotted line).

**Fig. S4.**
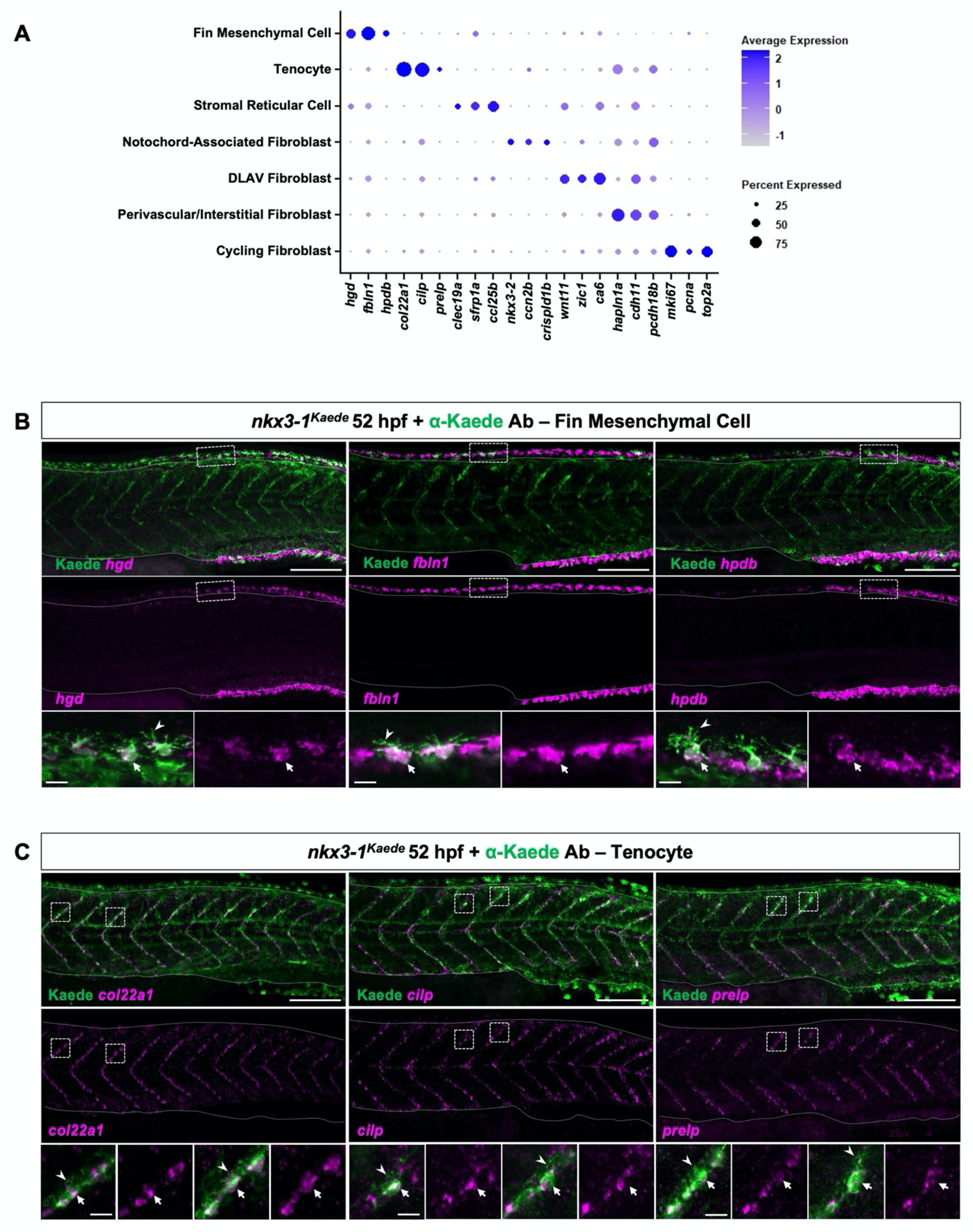
Top markers for fibroblast subclusters and mapping of fin mesenchymal cell and tenocyte clusters in the zebrafish trunk. Related to Fig. 1. (**A**) Expression profile of top 3 highly expressed markers for each fibroblast subtype identified in Fig. 1D. Coloration represents average expression level; circle size represents proportion of expressing cells per cluster. (**B,C**) Co-immunostaining (green, labelling *nkx3-1^Kaede^*-positive fibroblasts) and fluorescent in situ hybridization (magenta) in 52 hpf *nkx3-1^Kaede^* embryos, showing expression of fin mesenchymal cell (B) and tenocyte (C) marker genes in the zebrafish trunk. Fin mesenchymal cell markers (*hgd*, *Kln1*, *hpdb*) are expressed in fibroblasts (arrows) within the dorsal and ventral fin folds (B), while tenocyte markers (*col22a1*, *cilp*, *prelp*) label fibroblasts (arrows) along the myotendinous junctions (C). Notched arrowheads indicate characteristic processes of fin mesenchymal cells and tenocytes in B and C, respectively. Solid lines in B and C indicate dorsal and ventral edges of the trunk; dotted lines indicate location of insets. *n* = 15 embryos/probe. Scale bars: 100 μm (B,C); 10 μm (insets).

**Fig. S5.**
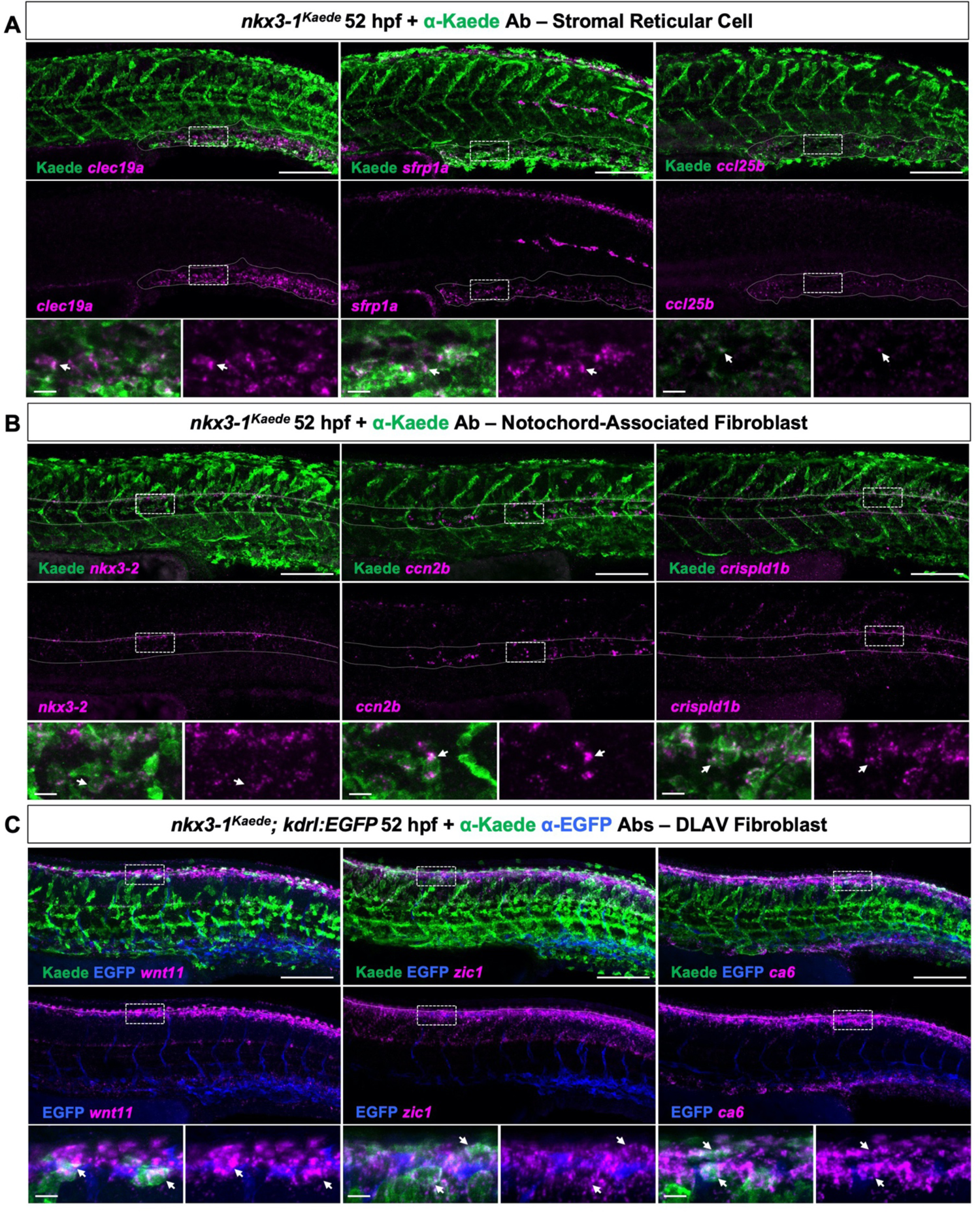
Distribution of stromal reticular cells, notochord-associated fibroblasts and DLAV fibroblasts in the zebrafish trunk. Related to Fig. 1. (**A-B**) Representative images of 52 hpf *nkx3-1^Kaede^*embryo trunks stained for sclerotome-derived fibroblasts (green, labelled by *nkx3-1^Kaede^*) and fibroblast subcluster marker genes (magenta) from Fig. S4A. (A) Stromal reticular cell markers (*clec19a*, *sfrp1a*, *ccl25b*) are expressed in fibroblasts (arrows) within the caudal vein plexus (solid lines). (B) Notochord-associated fibroblast markers (*nkx3-2*, *ccn2b*, *crispld1b*) are enriched in cells (arrows) surrounding the notochord (solid lines). (**C**) Fluorescent mRNA in situ hybridization for DLAV fibroblast markers (*wnt11*, *zic1*, *ca6*; magenta) in 52 hpf *nkx3-1^Kaede^; kdrl:EGFP* embryos, showing expression in fibroblasts (arrows, labelled by *nkx3-1^Kaede^*) around the dorsal longitudinal anastomotic vessel (DLAV) (blue, labelled by *kdrl:EGFP*) running along the dorsal edge of the trunk (solid lines). Dotted lines in A-C indicate location of inset. *n* = 15 embryos/probe. Scale bars: 100 μm (A-C); 10 μm (insets).

**Fig. S6.**
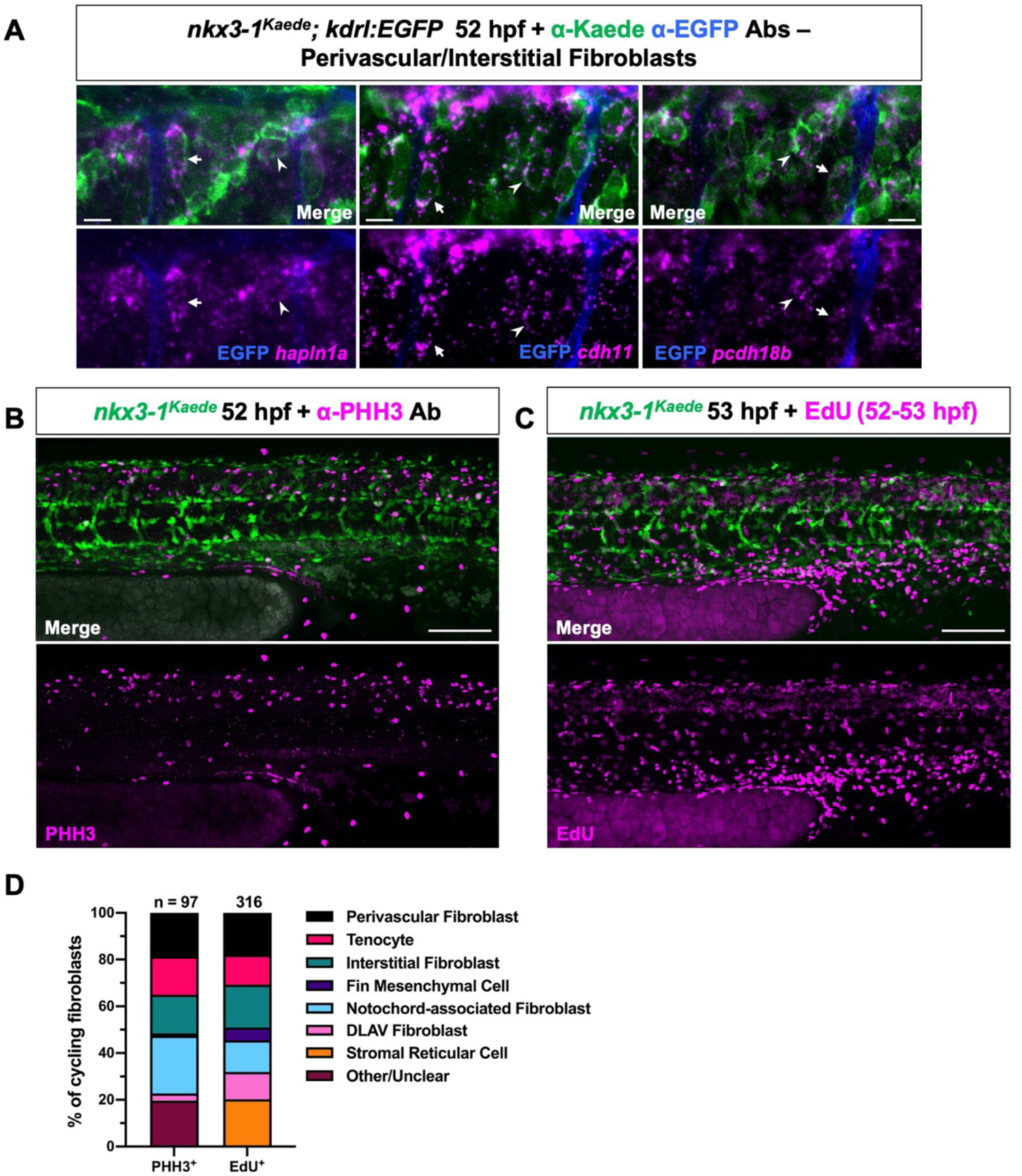
Mapping of perivascular/interstitial fibroblasts and cycling fibroblasts within the zebrafish trunk. Related to Fig. 1. (**A**) Fluorescent mRNA in situ hybridization in 52 hpf *nkx3-1^Kaede^; kdrl:EGFP* embryos showing expression of perivascular/interstitial fibroblast cluster markers (*hapln1a*, *cdh11*, *pcdh18b*) in interstitial fibroblasts (arrowheads) and perivascular fibroblasts (arrows) surrounding intersegmental vessels (blue, labelled by *kdrl:EGFP*). *n* = 15 embryos/probe. (**B**) Immunostaining for mitotic marker, phospho-histone H3 (PHH3) (magenta), in 52 hpf *nkx3-1^Kaede^* embryos. PHH3-positive fibroblasts are distributed throughout the embryonic trunk. *n* = 13 embryos. (**C**) Trunk region of 53 hpf *nkx3-1^Kaede^* embryos showing EdU incorporation (magenta) in fibroblasts (green) from 52 to 53 hpf. Proliferating EdU-positive fibroblasts can be seen throughout the trunk. *n* = 11 embryos. (**D**) Fibroblast subtype composition of the total cycling fibroblast populations from B,C. Subtype identity was determine based on morphology and location. *n* = 97 fibroblasts from 13 embryos (PHH3); 316 fibroblasts from 11 embryos (EdU). Scale bars: 10 μm (A); 100 μm (B,C).

**Fig. S7.**
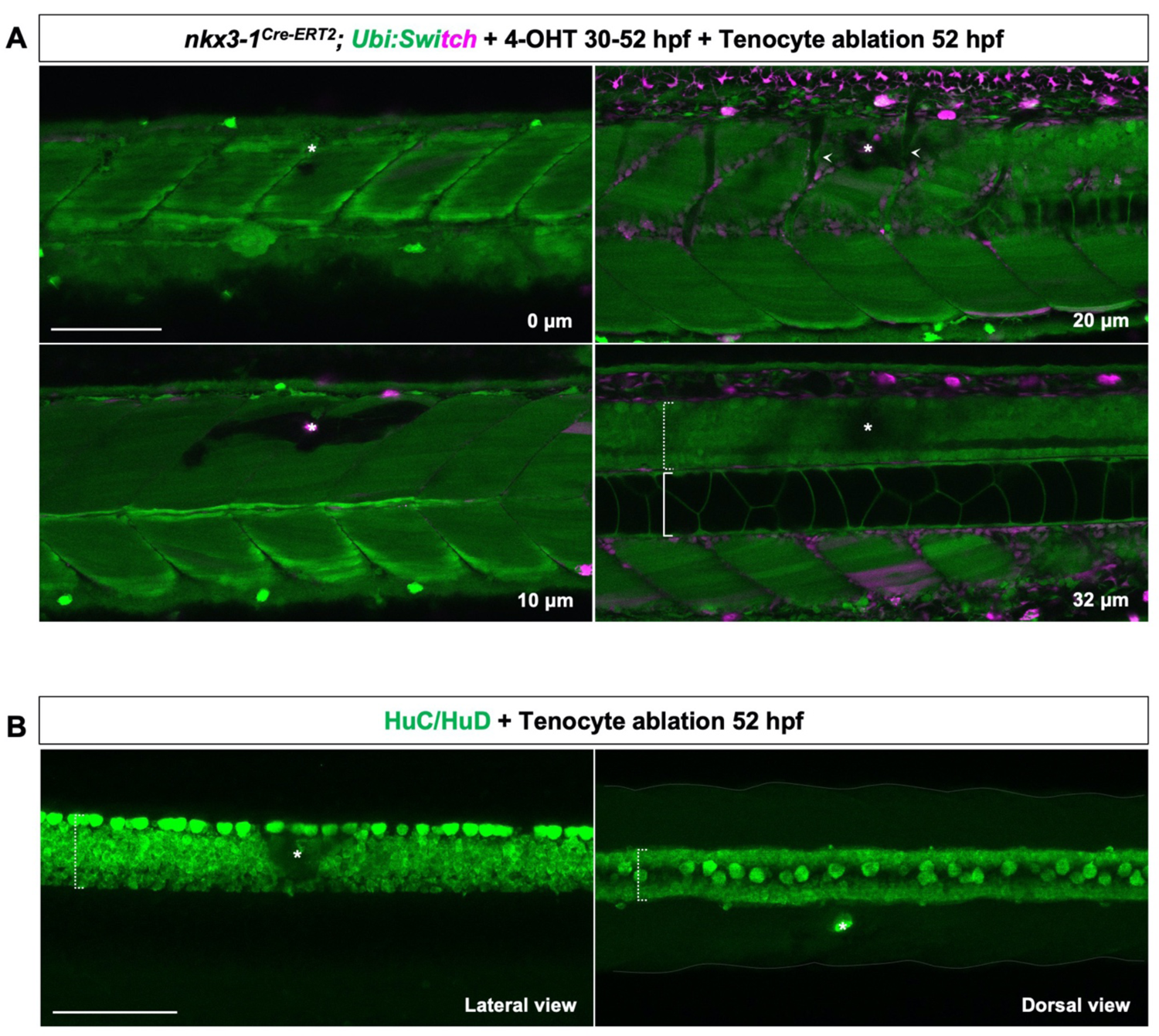
Laser ablation results in localized tissue loss. Related to Fig. 2. (**A**) Single z-slice images showing superficial and deeper tissues from the trunk region of an injured *nkx3-1^Cre-ERT2^; Ubi:Switch* embryo at 52 hpf, with slice depth shown. Embryos were treated with 4-OHT from 30 to 52 hpf prior to laser injury, resulting in the expression of mCherry in sclerotome-derived fibroblasts (magenta) and GFP in all other cells (green). Muscle fibers and the epithelium superficial to the injury site (asterisk) remained largely intact after laser ablation (0 μm slice). Ablation resulted in the loss of muscle fibers and mCherry^+^ fibroblasts surrounding the injury site (asterisks), but not ISVs (notched arrowheads) (10 and 20 μm slices). Laser ablation also led to the loss of fluorescent signal in the spinal cord (dotted bracket) immediately adjacent to the injury site although the notochord (solid bracket) was unaffected (32 μm slice). *n* = 12 embryos. (**B**) Lateral and dorsal views of the spinal cord of an injured embryo at 52 hpf, stained for neuronal marker HuC/HuD (green). While the lateral view (left) shows reduced staining in the spinal cord near the injury site (asterisks), the dorsal view (right) reveals that the spinal cord medial to the injury site is completely intact. This suggests that the loss of signal in lateral views (32 μm slice shown in A) is likely caused by blockage of the excitation/emission light by the opaque scar tissue, rather than spinal cord injury. Solid lines indicate the boundaries of the trunk. *n* = 15 embryos. Scale bars: 100 μm.

**Fig. S8.**
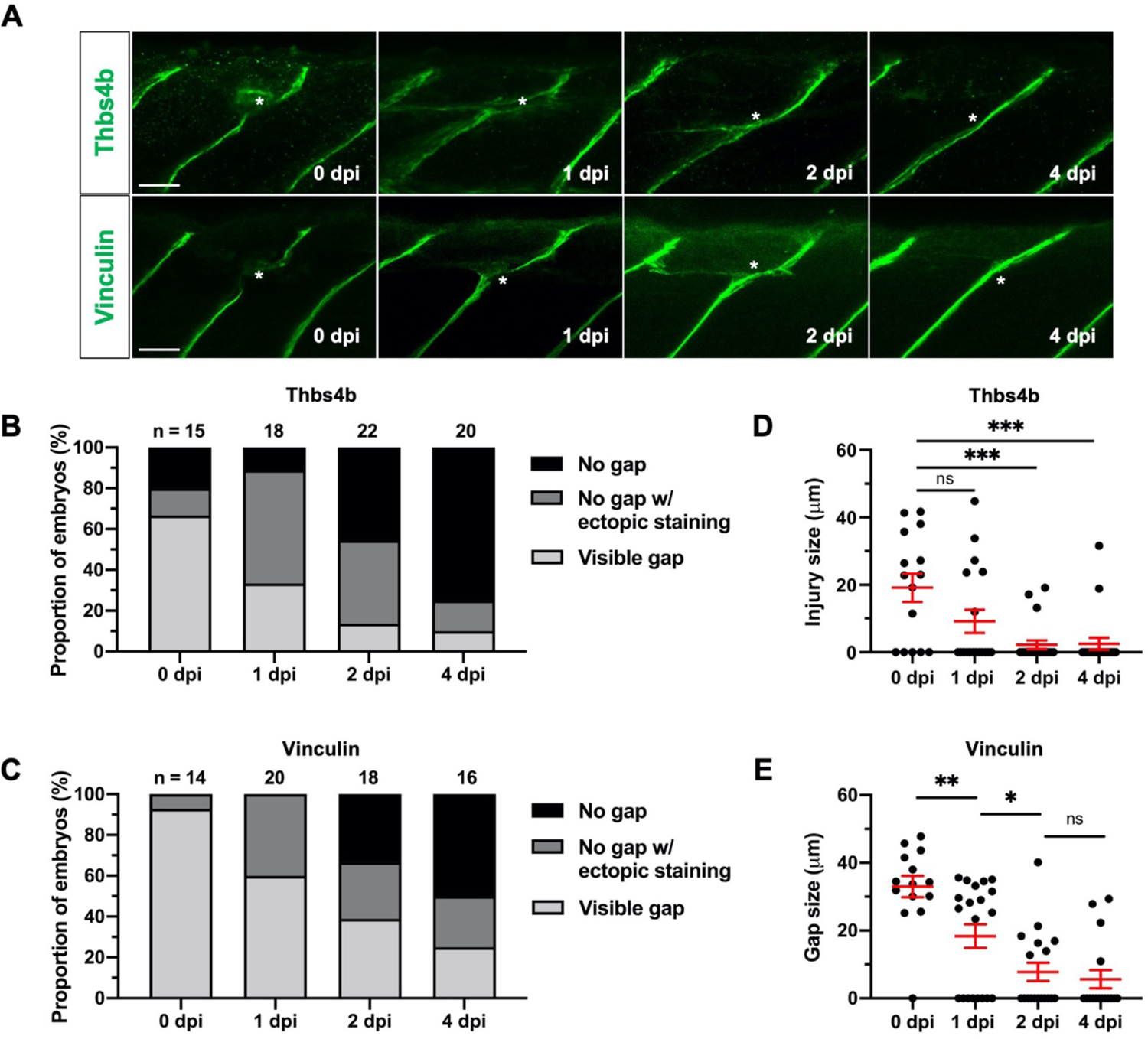
Zebrafish myotendinous junction (MTJ) architecture is restored after laser injury. Related to Fig. 2. (**A**) Immunostaining for MTJ markers, Thrombospondin 4b (Thbs4b) (top) and Vinculin (bottom), from 0 to 4 dpi. The MTJ extracellular matrix is restructured along the injury site (asterisks) following tenocyte ablation. *n* = 15-22 embryos/stage (Thbs4b); 14-20 embryos/stage (Vinculin). (**B,C**) Distribution of phenotypes in Thbs4b (B) and Vinculin (C) stained embryos from 0 to 4 dpi. Most injured embryos showed no obvious gap in staining by 4 dpi. (**D,E**) Quantification of gap in Thbs4b (D) and Vinculin (E) staining at the injured MTJ from 0 to 4 dpi. Data shown as mean ± SEM. Statistics: Mann-Whitney *U*-test. Significance: p-value >0.05 (ns), <0.05 (*), <0.01 (**), <0.001 (***). Scale bars: 25 μm.

**Fig. S9.**
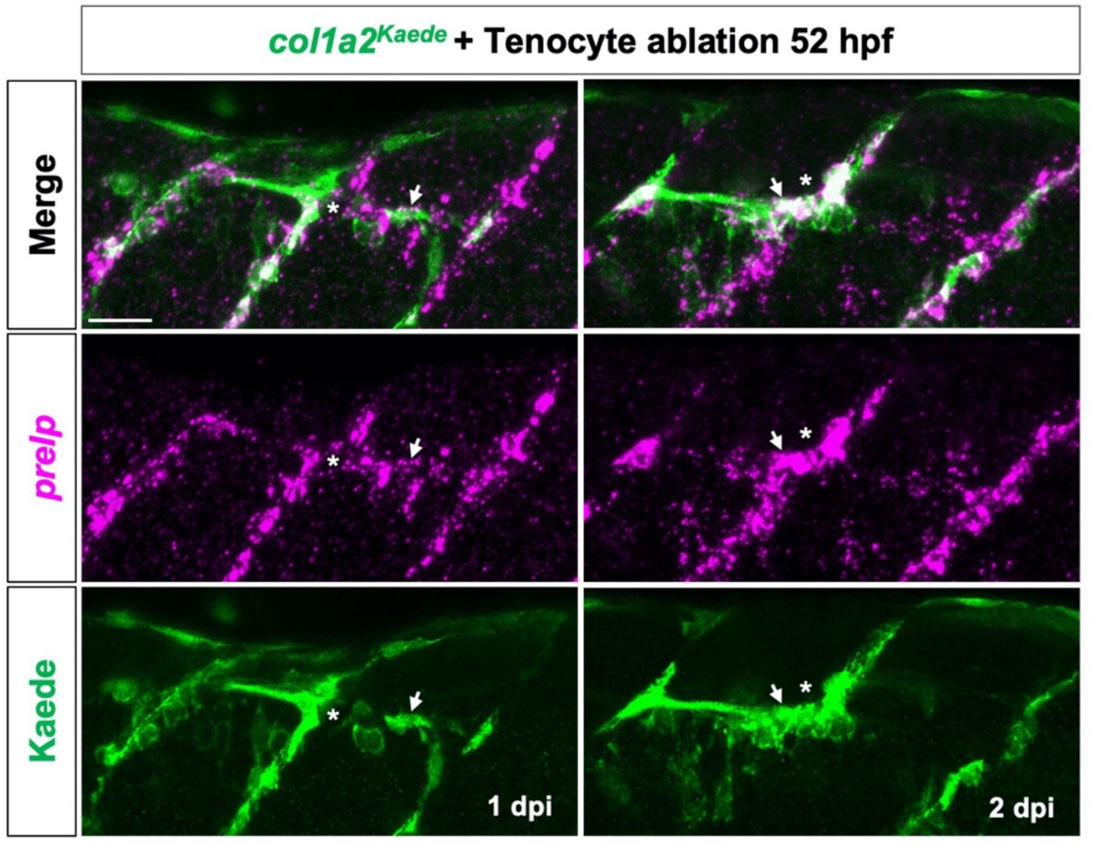
Injury-responsive fibroblasts show ectopic expression of tenocyte marker *prelp*. Related to Fig. 3. Fluorescent mRNA in situ hybridization showing expression of tenocyte marker *prelp* (magenta) in activated fibroblasts (arrows, labelled by *col1a2^Kaede^*) outside the MTJ at 1 and 2 dpi. Injury site is indicated by asterisk. *n* = 17 (1 dpi) and 14 (2 dpi) embryos. Scale bar: 25 μm.

**Fig. S10.**
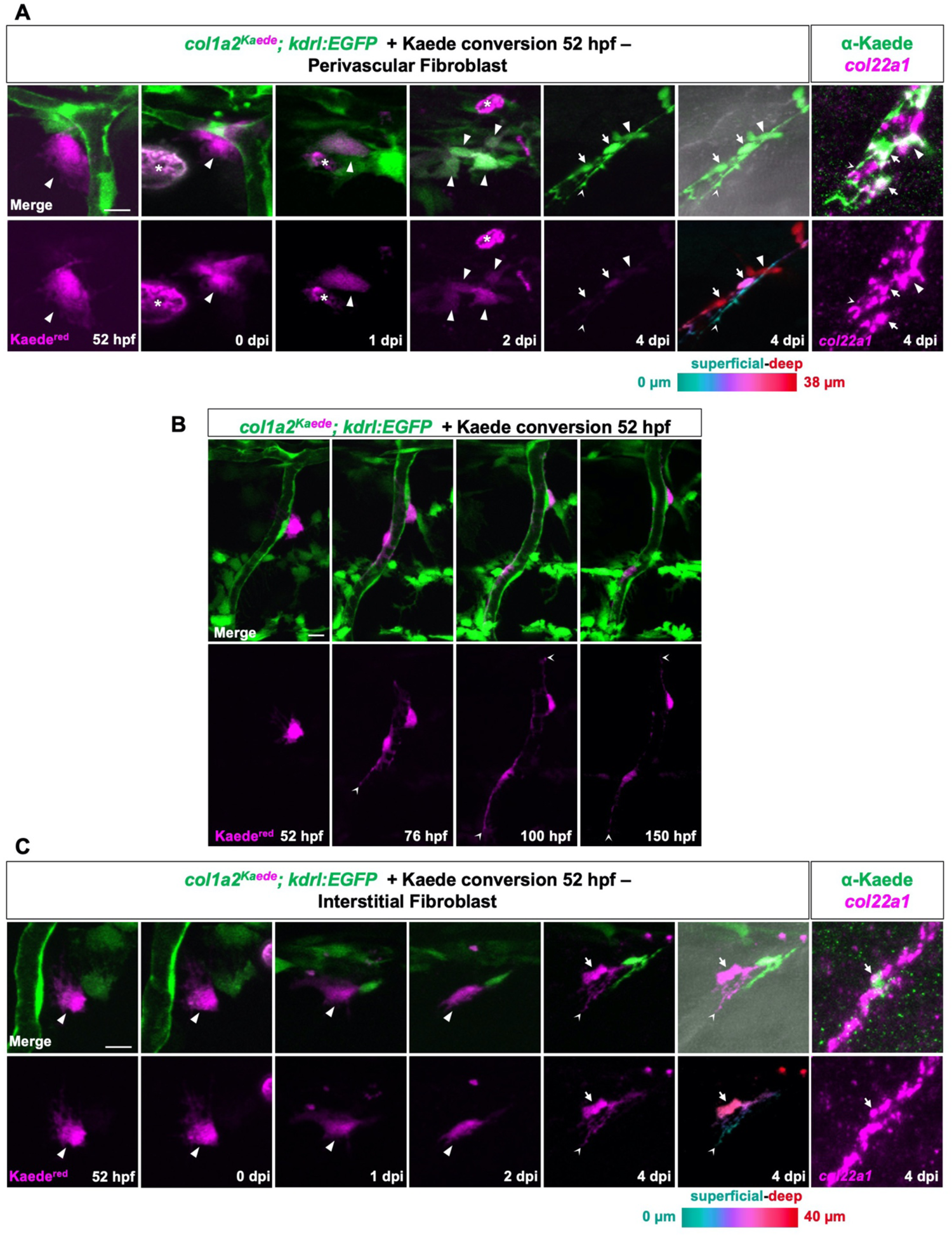
Perivascular and interstitial fibroblasts contribute to new tenocytes at the injury site. Related to Figs. 4 & 5. (**A**) Close-up views of a single photoconverted perivascular fibroblasts in an injured *col1a2^Kaede^; kdrl:EGFP* embryo from 0 to 4 dpi as shown in Fig. 4A. After fixation at 4 dpi, the same embryos were stained with α-Kaede Ab and tenocyte marker *col22a1*. In live images, the traced cell (arrowheads) can be seen gradually changing its globular morphology between 52 hpf and 2 dpi. By 4 dpi, some clonal progeny (arrows) show a stereotypical tenocyte morphology with small cell body and long superficially-extending processes (notched arrowheads), and express tenocyte marker *col22a1*. *n* = 21/22 cells from 16 embryos. (**B**) Images of an uninjured *col1a2^Kaede^; kdrl:EGFP* embryo from 52 to 150 hpf. The photoconverted perivascular fibroblast (magenta) divides once and its progeny can be seen transitioning toward a pericyte-like morphology with elongated cell bodies and long ISV-associated processes (notched arrowheads). *n* = 9 cells from 8 embryos. (**C**) Close-up images of a single photoconverted interstitial fibroblast in an injured *col1a2^Kaede^; kdrl:EGFP* embryo from 0 to 4 dpi. The same embryos were co-stained with α-Kaede Ab and tenocyte marker *col22a1* at 4 dpi. Like the perivascular fibroblast in A, the traced cell undergoes morphological changes and differentiates into a *col22a1*-positive tenocyte with long cellular processes (notched arrowheads). *n* = 6/6 cells from 6 embryos. Scale bars: 10 μm.

**Fig. S11.**
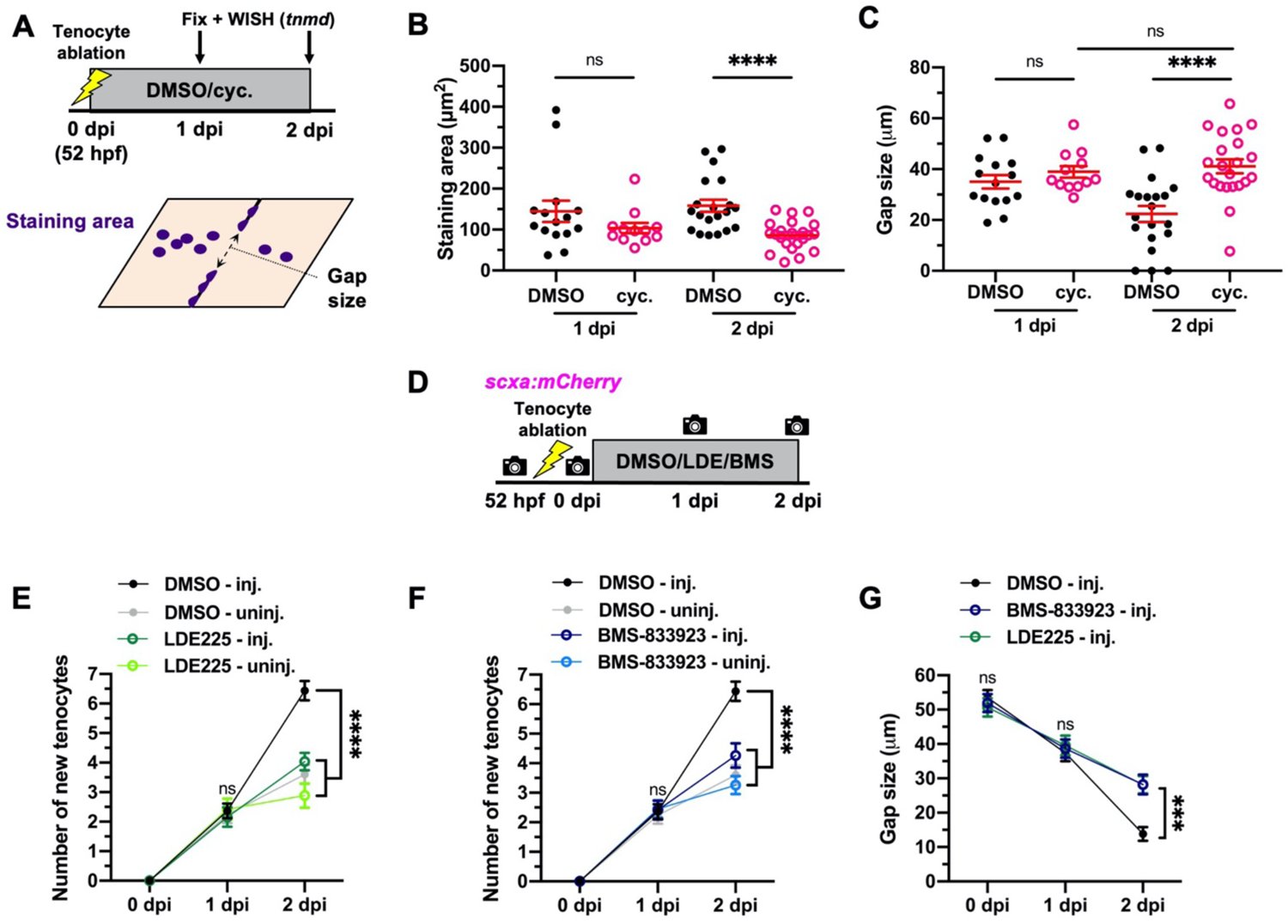
Inhibition of Hh signaling impairs tenocyte regeneration. Related to Fig. 7. (**A**) Drug treatment and staining protocol for embryos after tenocyte ablation. Tenocyte ablation was performed at 52 hpf and embryos were immediately transferred to DMSO or cyclopamine-containing fish water. Treatment was washed off prior to fixation and in situ hybridization was performed for tenocyte marker *tnmd*. Staining area and gap in staining were quantified as depicted. (**B,C**) Quantification of *tnmd* staining area (B) and gap size (C) around the injury site. Cyclopamine-treated embryos showed significantly lower staining area and larger gap between tenocytes at 2 dpi compared to DMSO-treated controls. *n* = 15 (1 dpi) and 20 (2 dpi) DMSO-treated embryos; 12 (1 dpi) and 22 (2dpi) cyclopamine-treated embryos. (**D**) Experimental protocol for tenocyte tracing in drug-treated *scxa:mCherry* transgenic embryos. Individual embryos were incubated in DMSO or Hh signaling inhibitors, LDE225 and BMS-833923, after injury and imaged repeatedly from 52 hpf to 2 dpi. Tenocytes numbers were manually recorded along injured and uninjured MTJs in each embryo. (**E,F**) Mean number of new tenocytes detected from 0-2 dpi in DMSO/LDE225 (E) or DMSO/BMS-833923 (F) treated embryos. Significantly fewer tenocytes were generated along injured MTJs in drug-treated embryos compared to DMSO-treated controls. (**G**) Mean gap in tenocytes at the injured MTJ in DMSO, BMS-833923, or LDE225-treated embryos from 0-2 dpi. *n* = 30 (injured) and 30 (uninjured) MTJs from 30 DMSO-treated embryos; 26 (injured) and 26 (uninjured) MTJs from 26 LDE225-treated embryos; 29 (injured) and 29 (uninjured) MTJs from 29 BMS-833923-treated embryos. Data are plotted as mean ± SEM. Statistics: Mann-Whitney *U* test (B,C); Sidak’s multiple comparisons (E-G). Significance: p-value >0.05 (ns), <0.001 (***), <0.0001 (****).

**Fig. S12.**
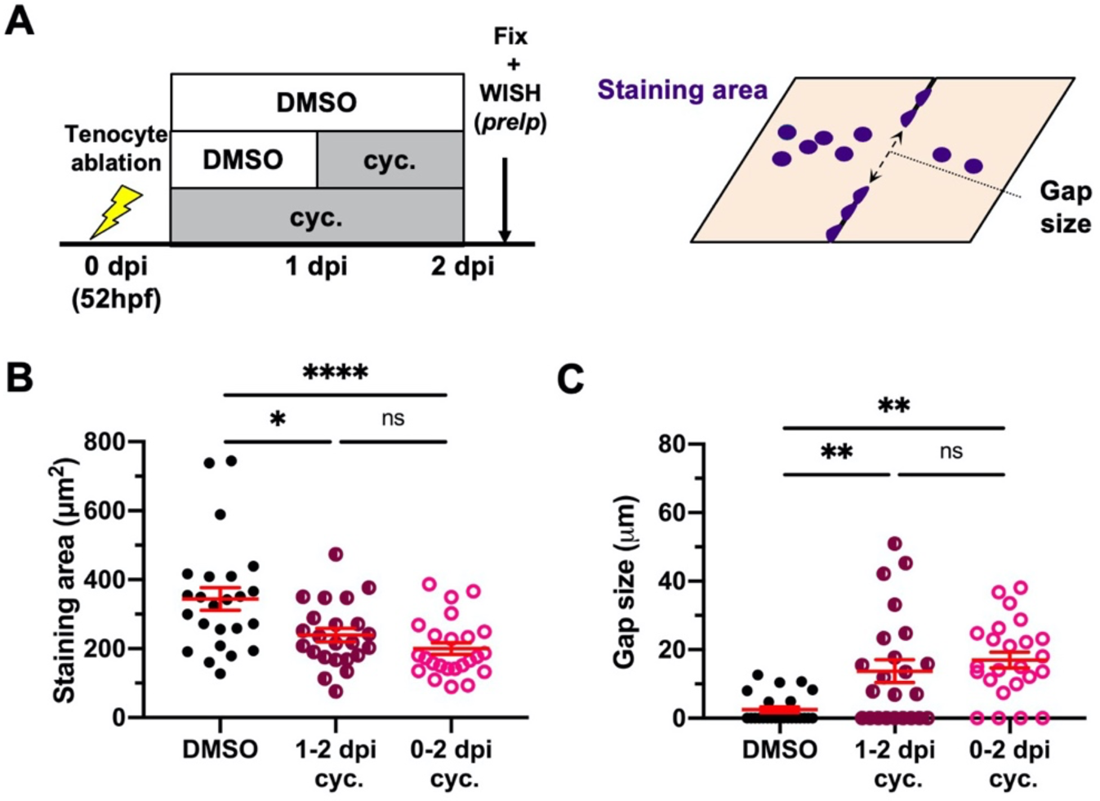
Short-term inhibition of Hh signaling compromises tenocyte regeneration. Related to Fig. 7. (**A**) Experimental protocol to test effectiveness of short-term cyclopamine treatment on tenocyte regeneration. Injured embryos were treated with DMSO or cyclopamine from 0-2 dpi or 1-2 dpi, prior to fixation and in situ hybridization for tenocyte marker *prelp*. Staining area and gap size were quantified as depicted. (**B,C**) Quantifications of *prelp* staining area (B) and gap size (C) surrounding the injury site. Both short and long-term cyclopamine treatment resulted in significantly larger gap size and reduced staining area compared to DMSO-treated controls. *n* = 24 DMSO-treated embryos; 23 (1-2 dpi) and 24 (0-2 dpi) cyclopamine-treated embryos. Data are plotted as mean ± SEM. Statistics: Mann-Whitney *U* test. Significance: p-value >0.05 (ns), <0.05 (*), <0.01 (**), 0.0001 (****).

**Table S1. Single-cell RNA sequencing cluster markers. Related to Figs. 1B & S2.** All markers identified for each cluster were obtained from scRNA-seq of mCherry^+^ cells as shown in Fig. 1B. Markers were selected using the Seurat ‘FindAllMarkers’ function with following specified parameters: logfc.threshold = 0.25.

**Table S2. Top markers of sclerotome-derived fibroblasts. Related to Figs. 1D & S4A.** Highly expressed marker genes are listed for subsetted sclerotome-derived fibroblast clusters as shown in Fig. 1D. Markers were selected using the Seurat ‘FindAllMarkers’ function with the following specified parameters: logfc.threshold = 0.25, min.diff.pct = 0.3.

**Table S3.**
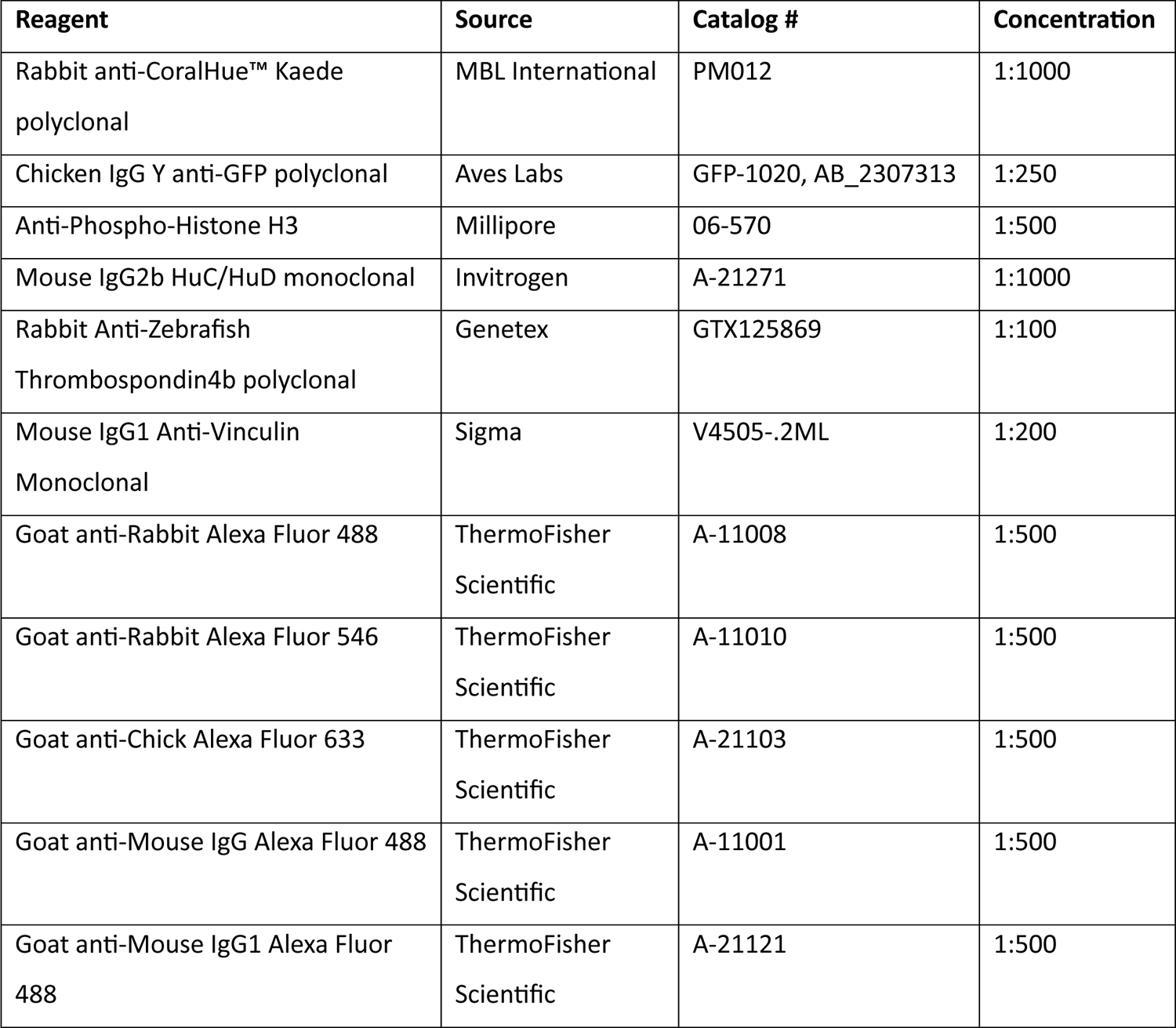
List of antibodies used in the present study.

**Movie S1. Perivascular fibroblasts respond to tendon injury.** Time-lapse imaging of a single photoconverted perivascular fibroblast (arrowhead) in an injured *col1a2^Kaede^; kdrl:EGFP* embryo from 0 to 24 hours post tenocyte ablation. The traced cell can be seen extending processes, migrating from the intersegmental vessel to the injury site (asterisk), and undergoing one round of cell division in the recorded period. Embryos were imaged at 20-minute time intervals. Time indicated in hh:mm format. *n* = 10 embryos. Scale bars: 50 μm.

